# Local differences in baseline sodium shape astrocytic potassium uptake by the NKA

**DOI:** 10.1101/2025.11.18.687951

**Authors:** Jan Meyer, Sara Eitelmann, Alok Bhattarai, Viola Bornemann, Petr Unichenko, Simone Durry, Karl W. Kafitz, Christian Henneberger, Ghanim Ullah, Christine R. Rose

## Abstract

Astrocytes are vital for the maintenance of ion and transmitter homeostasis in the extracellular space, with the inward Na⁺ gradient playing a pivotal role in these processes. Earlier studies not only reported a low baseline Na^+^ concentration ([Na^+^]) in astrocytes, but also suggested an equilibration of [Na^+^] within the gap-junction-coupled syncytium. This is consistent with the view that the basic homeostatic properties of astrocytes are largely identical due to their critical role in brain function. Here, we used multi-photon fluorescence lifetime imaging for a quantitative determination of astrocytic [Na^+^] in mouse forebrain tissue slices and *in vivo*. Contrary to the prevailing notion of a rather uniform Na⁺ distribution, we detected a previously unobserved subcellular and cellular heterogeneity in astrocytic [Na^+^], accompanied by differences in the capacity for Na^+^/K^+^-ATPase (NKA)-mediated uptake of extracellular K^+^. Biophysical modelling showed that this heterogeneity can be replicated by the reported differential expression of NKA isoforms in astrocytes together with a different strength of Na^+^ influx over the plasma membranes. Altogether, our results thus suggest the existence of functionally distinct astrocytes and astrocyte subdomains in which Na^+^ homeostasis is locally adapted to the specific requirements of surrounding neural networks.

## Introduction

Astrocytes are essential for the proper functioning of the vertebrate brain. They contribute to the formation and plasticity of neural networks ^1–3^, and are critically involved in the ionic homeostasis of the extracellular space (ECS) ^4,5^. This includes the uptake of K^+^ by Kir4.1 channels and by the Na^+^/K^+^-ATPase (NKA) ^6–9^. The control of extracellular K^+^ ([K^+^]_e_) is a central mechanism by which astrocytes control neuronal excitability and network performance ^10^. In addition to its well-known role in [K^+^]_e_ homeostasis, the NKA also represents the dominant mechanism for the export of Na^+^, establishing a low intracellular Na^+^ concentration ([Na^+^]) against a high inward Na^+^ gradient ^11,12^. The latter provides the driving force for a multitude of plasma membrane transporters including Na^+^/K^+^/2Cl^-^ cotransporters (NKCC1), Na^+^/H^+^ exchangers (NHE) or Na^+^-dependent glutamate transporters (EAATs). These challenge the Na^+^ gradient both at rest and during neuronal activity, making a low [Na^+^] and NKA activity the “core and hub” of astrocyte function ^13–15^.

Mean values for [Na^+^] determined in astrocytes from rodent brain tissue slices are in the range of 12-15 mM ^14,16,17^. This data, however, represents bulk measurements from cell bodies, and it is currently unclear whether somatic [Na^+^] ([Na^+^]_s_) also reflects [Na^+^] in glial processes ([Na^+^]_p_), or whether the latter exhibit a different [Na^+^]. Due to the high mobility of Na⁺ and the absence of relevant Na⁺-buffers, it is commonly assumed that at rest, [Na⁺] is largely equilibrated within individual astrocytes and between gap-junction-coupled glial syncytia ^18–21^. Since a low [Na⁺] is crucial for ionic homeostasis, it also seems obvious that baseline [Na⁺] should be essentially identical in different astrocytes and astrocyte sub-compartments to ensure the reliable fulfillment of vital astrocytic functions.

Recent modeling work, however, challenged this concept, proposing the existence of microdomains and subcellular gradients for Na^+^ ^22,23^. Along this line, large local depolarizations mediated by neuronal activity and activation of EAATs were reported from peripheral astrocyte processes, indicating the existence of local (Na^+^-) signaling domains ^24,25^. Moreover, the efficacy of local (Na^+^-dependent) glutamate uptake by astrocytes depends on the spine size ^26^ as well as on the spatial proximity of an astrocyte process to a synapse ^27^, again pointing towards a likely subcellular heterogeneity of Na^+^ influx.

As mentioned above, there is currently no experimental data addressing potential heterogeneities in astrocytic [Na⁺]. One reason for this lack is that the unbiased determination of astrocytic [Na⁺] is challenging. At present, there are no genetically-encoded Na⁺ indicators suitable for use within neural cells in brain tissue. Moreover, intensity-based measurements using chemical indicators are prone to artefacts related to differences in, or changes of, dye concentrations. An alternative approach, however, is fluorescence lifetime imaging microscopy (FLIM), which is essentially independent on dye concentrations ^28^, and has for example enabled quantitative analysis of astrocytic Ca^2+^ or Cl^-^ concentrations ^9,29–32^.

Here, we established multi-photon fluorescence lifetime imaging (MP-FLIM) based on time-correlated single-photon counting (TCSPC) of the Na^+^ indicator ION Natrium Green-2 (ING-2) ^33,34^ for quantitative, dynamic imaging of astrocytic [Na^+^] in mouse forebrain tissue slices and *in vivo*. Overall, our results reveal a hitherto unexpected cellular and subcellular heterogeneity in astrocytic Na^+^-homeostasis, resulting in functionally distinct astrocyte subgroups and subdomains, optimized for efficient regulation of ion and transmitter homeostasis in the ECS.

## Results

### MP-FLIM-based measurement of astrocytic [Na^+^]_s_ *in situ* and *in vivo*

So far, FLIM has been employed for measurement of astrocytic [Ca^2+^] or [Cl^-^] in brain tissue slices and *in vivo* ^9,29–32^. To establish MP-FLIM of astrocytic [Na^+^]_s_, hippocampal slices were bolus-loaded with ING-2 (Fig. 1a), resulting in the staining of SR101-positive astrocytes in the CA1 *stratum radiatum*. This enabled the recording of the fluorescence lifetime (FL) of ING-2 in regions of interest (ROIs) placed around astrocyte somata using the SR101 fluorescence channel (Fig. 1b). Slices were then perfused with different calibration salines containing a defined [Na^+^] (0-150 mM) (Fig. 1c). ING-2 FL increased with increasing [Na^+^]_s_, following a Michaelis-Menten relationship with an apparent K_m_ of 21 mM, demonstrating that MP-FLIM of ING-2 is well suited for determination of astrocytic [Na^+^] ^33^ (Fig. 1c-e).

**Fig. 1:**
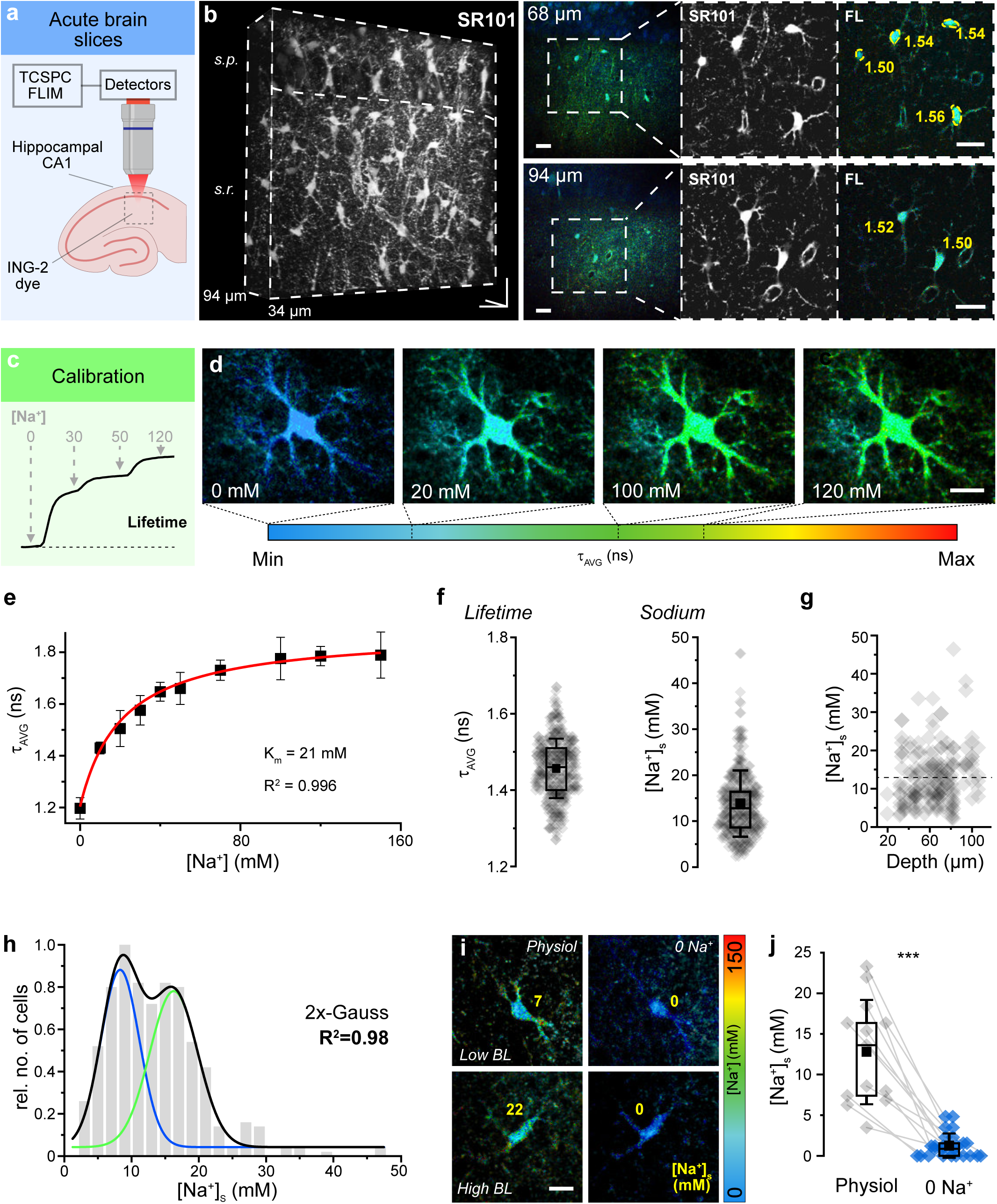
MP-FLIM-based determination of astrocyte [Na^+^] in the hippocampal CA1 area. **a** Scheme of experimental design. **b** Z-Stacks of SR101 fluorescence intensity and ING-2 FL. *Left*: 3D reconstruction of SR101 (depth: 34-94 μm). *s.p.: stratum pyramidale*, *s.r.*: *stratum radiatum*, Scale: 20x20x60 μm. *Right:* Two optical planes (depth: 68 and 94 µm) from the *s.r.* of the same slice. 1^st^ column: ING-2 FL, 2^nd^ column: SR101 intensity, 3^rd^ column: ING-2 FL of the boxed region with an SR101 mask employed (for detailed description, see Methods and Suppl. Fig. S2). Top right: ROIs are indicated exemplarily in yellow. Numbers indicate the FL determined from somata. Scales: 20 μm. **c** Scheme of calibration strategy, indicating changes in FL with changes in [Na^+^]. **d** Color-coded FL images (masked with SR101) of a single, bolus-stained astrocyte exposed to calibration salines containing the indicated [Na^+^]. Color-code, showing the average, amplitude-weighted FL (τ_AVG_) is depicted below. Scale: 10 μm. **e** Relation between τ_AVG_ and [Na^+^]. Red line: Michaelis-Menten fit. Note that FL saturates at around 1.8 ns/100 mM Na^+^. **f** τ_AVG_ (left) and somatic [Na^+^]_s_ (right) of astrocytes. **g** Correlation between [Na^+^]_s_ and depth of the optical section (Pearson: 0.17, R^2^=0.03). **h**[Na^+^]_s_. Black line represents a double Gaussian function, composed of two individual Gaussian distributions (blue and green line). **e-h**: n=369, N=32. **i** FL images (masked with SR101) of two astrocytes with initially low (top) and initially high (bottom) baseline [Na^+^]_s_ (Low/High BL) in control (Physiol) and after perfusion with Na^+^-free calibration saline (0 Na^+^). Yellow numbers indicate [Na^+^]_s_. Scale: 10 µm **j** [Na^+^]_s_ of astrocytes in control (Physiol) and in Na^+^-free calibration solution (0 Na^+^). Data points from individual cells are connected by lines (n=18, N=4; Wilcoxon signed rank test, p=2.14092E-4). **e, f, g, j:** diamonds: individual data points, boxes: 25/75, whiskers: SD, lines: median, squares: mean.

To determine astrocytic baseline [Na^+^]_s_, ING-2 FL was determined in somata of astrocytes in slices perfused with standard saline (Fig. 1f, left). Converting this data using the *in situ* calibration curve revealed a mean [Na^+^]_s_ of 13.8±7.2 mM (median: 12.8 mM; n=369, N=32) (Fig. 1f, right). The vast majority of cells (80%) exhibited a [Na^+^]_s_ between 5 and 20 mM (total range: 2-46 mM). [Na^+^]_s_ was independent on the relative depth, and cells with high and low baseline [Na^+^]_s_ were both found close to the slice surface as well as in deeper tissue layers (Pearson 0.17, R^2^=0.03) (Fig. 1g). The overall distribution of astrocytic [Na^+^]_s_ could be best described by a double Gaussian fit (R²=0.98), centered around 8.2±0.5 mM and 16.3±0.7 mM (Fig. 1h).

In a further set of experiments, the FL of astrocytes was first determined in standard saline, revealing a mean baseline [Na⁺]_s_ of 12.8±6.4 mM (n=18, N=4) (Fig. 1 i,j). Afterwards, slices were perfused with nominally Na^+^ free calibration saline. Independent of their former baseline, washout of Na^+^ caused the FL in astrocyte somata to drop to values corresponding near 0 mM [Na⁺]_s_ (mean 1.3±1.5 mM), confirming that ING-2 FL levels determined in standard ACSF are reliable reporters of astrocytic [Na⁺]_i_ (Fig. 1 i-j).

Astrocyte properties differ among brain regions ^35–38^. To reveal potential differences between hippocampal and cortical astrocytes, we performed MP-FLIM in layers 2/3 of cortical slices (Fig. 2 a-d). Average [Na^+^]_s_ of cortical astrocytes was similar to that determined in hippocampal astrocytes (mean: 12.5±4.2 mM, median: 12 mM, range: 6.3-29.6 mM; n=34, N=4; p=0.55) (Fig. 2c). As observed in the hippocampus, [Na^+^]_s_ of cortical astrocytes was independent on the tissue depth (Pearson 0.04, R^2^=0.002) (Fig. 2d).

**Fig. 2:**
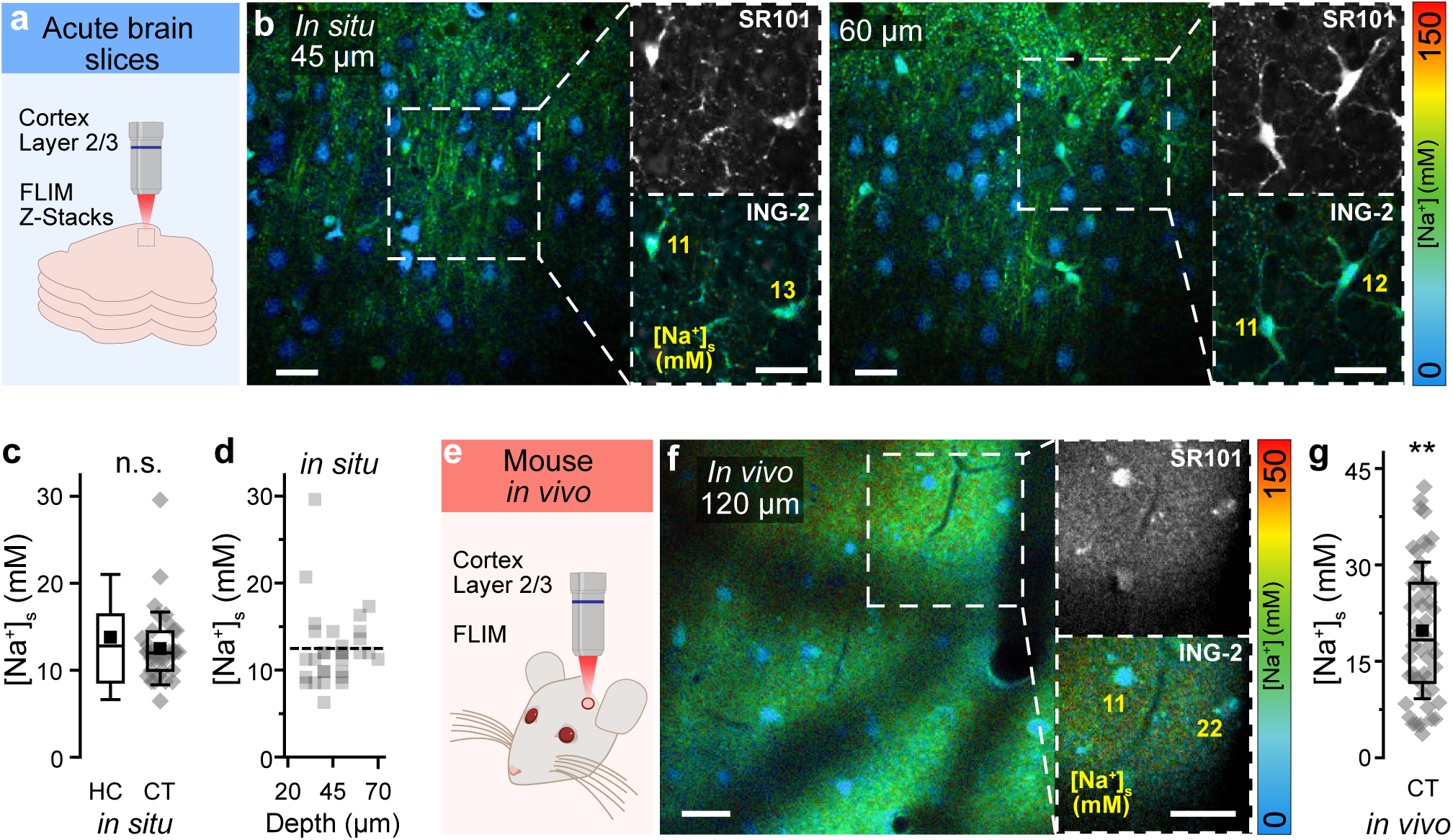
[Na^+^] of cortical astrocytes *in situ* and *in vivo*. **a** Scheme of experimental design of MP-FLIM in cortical tissue slices. **b** FL images of a cortical tissue slice at 45 µm (left) and 60 µm (right) of depth. Smaller images on the right depict the SR101 channel (top) and the ING-2 FL (masked with SR101) of the boxed regions at higher magnification. **c** [Na^+^]_s_ in hippocampal (HC, data taken from Fig. 1f) and cortical slices (CT; n=34, N=4; Mann-Whitney-Test: p= 0.55). **d** Correlation between [Na^+^]_s_ of cortical astrocytes and the depth of the optical section (Pearson: 0.17, R^2^=0.03). **e** Scheme of experimental design of MP-FLIM performed *in vivo*. **f** FL image at 120 µm of depth. Boxed regions on the right show SR101 and color-coded ING-2 FL (masked with SR101) at higher magnification. **g** Astrocytic [Na^+^]_s_ *in vivo* (n=50, 12 field of views; Mann-Whitney-Test vs. CT *in situ*: p=0.0017). **b, f:** Color-code for ING-2 FL is shown on the right. Yellow numbers indicate [Na^+^]_s_ in the indicated cells. **c, g:** diamonds: individual data points, boxes: 25/75, whiskers: SD, lines: median, squares: mean. All scales 20 μm.

Finally, we also assessed [Na^+^]_s_ *in vivo* performing MP-FLIM of ING-2 in SR101-positive cells of cortical layers 2/3 of anesthetized mice (Fig. 2 e-g). In this set of experiments, FL was converted into [Na^+^] by adapting an *in vitro* calibration to fit *in vivo* measurement conditions (see methods). Average astrocytic [Na^+^]_s_ was 19.8±10.6 mM (median: 18.3 mM, range: 3.7-42.1 mM; n=50, 12 field of views), which is significantly higher than in tissue slices of cortex (p=0.002) and hippocampus (p=0.001) (Fig. 2g).

Taken together, these results demonstrate that MP-FLIM of ING-2 enables determination of astrocytic [Na^+^]_s_ *in situ* and *in vivo*. While our experiments confirm earlier work by showing that mean somatic [Na^+^]_s_ is around 14 mM in hippocampal slices, they also reveal that baseline [Na^+^]_s_ of hippocampal astrocytes shows a rather large intercellular heterogeneity and displays a bimodal distribution centered around 8 and 16 mM. Moreover, we found that average baseline [Na^+^]_s_ is similar in hippocampal and cortical astrocytes *in situ*, but increased by several mM in cortical astrocytes *in vivo*.

### [Na^+^] in astrocyte processes

For quantification of [Na^+^] in astrocytic processes ([Na^+^]_p_), we loaded individual SR101-positive cells in the CA1 *stratum radiatum* with ING-2 using whole-cell patch-clamp (Fig. 3a, left). Cells exhibited a resting membrane potential of -84.8±0.4 mV and a linear I/V relationship, typical for mature astrocytes (n=24, N=24; not illustrated). Z-stacks of ING-2 FL were taken (step size 0.5 µm), covering a depth of at least 50 µm (frame binning of ≥10) (Fig. 3a, right). Somata of cells held in whole-cell configuration were excluded from FL analysis (Fig. 3a, right), because somata are quickly dialyzed by the intracellular saline, preventing a meaningful conversion of ING-2 FL into [Na^+^]_s_ ^34^.

**Fig. 3:**
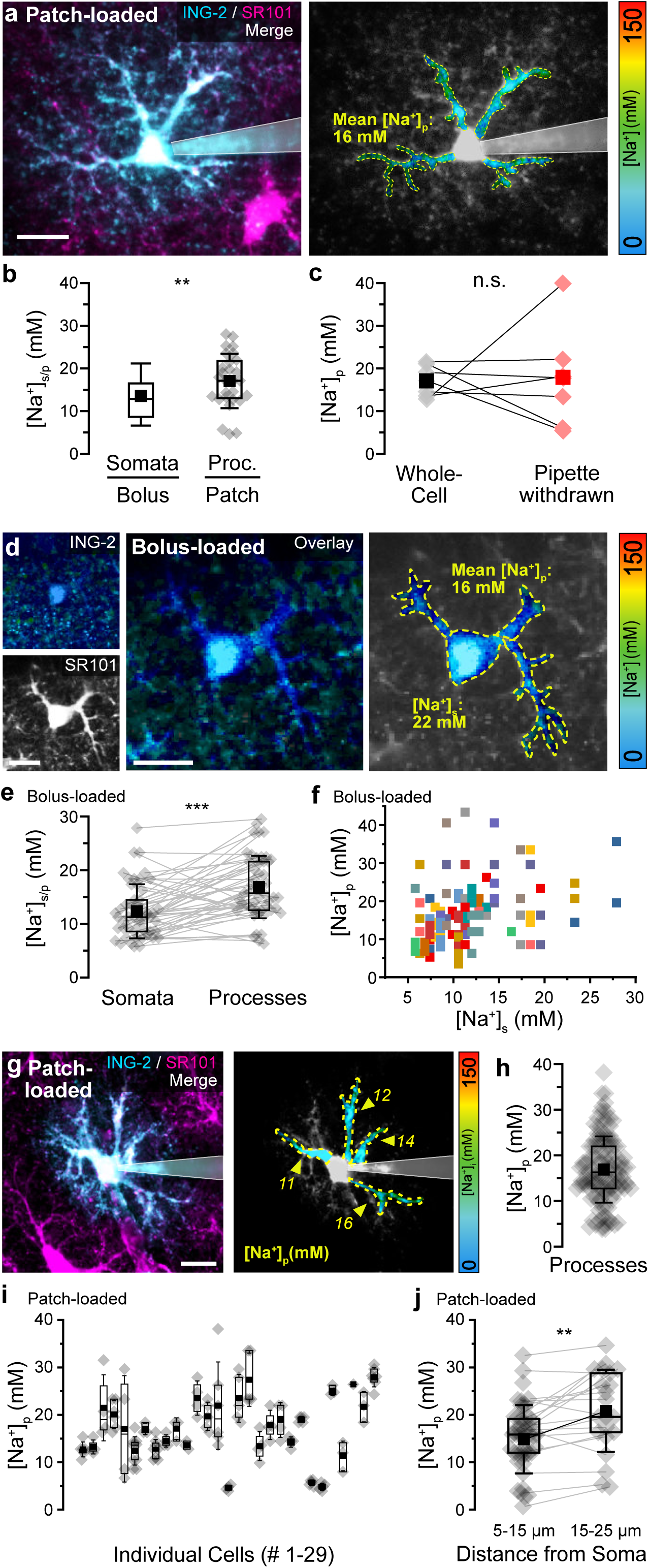
[Na^+^] in processes of hippocampal astrocytes *in situ*. **a** Left: Merge of ING-2 (cyan) and SR101 (magenta) intensity images of an astrocyte loaded with ING-2 via whole-cell patch-clamp. Right: color-coded ING-2 FL image. Colored, encircled regions indicate ROIs from which FL was determined. Note that the soma was excluded from analysis. **b** [Na^+^]_s_ in bolus-loaded slices (data taken from Fig. 1f) and average [Na^+^]_p_ per cell determined from patch-clamped astrocytes (n=28 cells, N=28; Mann-Whitney-Test, p=0.0018). **c** Average [Na^+^]_p_ per cell determined in whole-cell mode and after withdrawal of the patch pipette (n=7, N=7; Paired-sample-t test, p=0.93). Lines connect data points from individual cells. **d** Left: Images of ING-FL and SR101 fluorescence of a bolus-loaded astrocyte. Center: ING-2 FL image masked with SR101. Right: Color-coded image illustrating [Na^+^]_p_ and [Na^+^]_s_. Regions from which FL was determined are highlighted in color and are encircled by dotted lines. **e** Paired [Na^+^]_p_ and [Na^+^]_s_ data determined from bolus-loaded astrocytes (n=50, N=5; Wilcoxon-signed-rank test, p=0.0004). **f** Relation between [Na^+^]_s_ and [Na^+^]_p_ (n=50, N=5; Pearson: 0.4, R^2^=0.11). Identical colors in the same column indicate processes from the same astrocyte. **g** Left: Merge of ING-2 (cyan) and SR101 (magenta) intensity images of an astrocyte loaded via whole-cell patch-clamp. Right: color-coded ING-2 FL image. Dotted lines encircle processes from which FL was determined individually. Note that the soma was excluded from analysis. **h** [Na^+^]_p_ as determined in individual processes (161 processes, n=41, N=29). **i** [Na^+^]_p_ depicted for each individual cell (#1-29). **j** [Na^+^]_p_ derived from regions of interest at 5-15 (n=51, N=20) and 15-25 µm (n=21, N=14) from the soma. Lines connect data points derived from the same process (Two-sample-t test, different var., p=0.015). **a, d, g:** Color-code for ING-2 FL is shown on the right. Yellow numbers indicate [Na^+^] in the indicated ROIs. **b, c, e, h, i, j:** diamonds: individual data points, boxes: 25/75, whiskers: SD, lines: median, squares: mean. All scales: 20 μm.

ROIs were drawn around individual processes arising from the soma including clearly visible branchpoints and averaged for a given cell (Fig. 3a, right). This resulted in a mean baseline [Na^+^]_p_ of 17.3±6.5 mM (median: 17.1 mM, range: 5-27 mM; n=29 cells, N=29), which is significantly higher than [Na^+^]_s_ determined for somata in bolus-loaded slices (p=0.004) (Fig. 3b). Average [Na^+^]_p_ of a given cell was not correlated with its resting membrane potential (n=24, N=24; Pearson: -0.11, p=0.59; not shown). We next tested if dialysis by the pipette saline influenced [Na^+^]_p_. To this end, [Na^+^]_p_ was first determined with cells held in whole-cell mode, before the patch-pipette was gently withdrawn. After cells were allowed to reseal for at least 15 minutes, average [Na^+^]_p_ was not different from that in whole-cell mode (whole-cell: 17.0±3.6 mM, median: 17.1 mM; pipette withdrawn: mean: 17.5±11.7 mM, median: 17.7 mM; n=7, N=7; p=0.92) (Fig. 3c). This result shows that [Na^+^]_p_ is apparently not clamped by the pipette saline ([Na^+^]: 11.2 mM), again similar to what was observed before in primary dendrites of CA1 neurons ^34^.

As an alternative approach for determination of [Na^+^]_p_, we re-visited experiments performed before in bolus-loaded slices. After drawing ROIs around processes of individual cells using images of the SR101-fluorescence, ING-2 FL was evaluated from the same focal plane (Fig. 3d). Average [Na^+^]_p_ of single cells determined in bolus-loaded slices was 16.9±5.8 (median: 15.7, range: 6-30 mM; n=50, N=6). This is significantly higher than [Na^+^]_s_ determined in the same set of astrocytes (12.4±5.1 mM, median: 11.3 mM, range: 6-28 mM; p=5.34E-06) (Fig. 3e), but similar to average [Na^+^]_p_ of cells dye-loaded by patch-clamp (see above; p=0.81).

Moreover, we found that in bolus-loaded astrocytes, [Na^+^]_p_ in individual processes of a given cell was weakly linearly correlated with [Na^+^]_s_ (n=50, N=5; Pearson: 0.4, p=0.0038, R^2^=0.11) (Fig. 3f).

Next, we studied the variability of [Na^+^]_p_ within a given astrocyte dye-loaded via a patch-pipette (Fig. 3g). Averaging 131 individual processes of 29 cells held in whole-cell mode resulted in a mean [Na^+^]_p_ of 16.9±7.2 mM (Fig. 3h). Within the same astrocyte, [Na^+^]_p_ varied on average by 6.6±5.9 mM between individual branches, differing by as much as 25 mM (Fig. 3i). Moreover, we found that [Na^+^]_p_ was dependent on the distance from the soma. At a distance between 5-15 µm, average [Na^+^]_p_ was 14.9±7.2 mM (n=51, N=20), while at 15-25 µm, average [Na^+^]_p_ increased to 20.8±8.7 mM (n=21, N=14; p=0.005) (Fig. 3j).

Altogether, these results demonstrate that the [Na^+^]_p_ of a given astrocyte is significantly higher than that in the soma and increases with increasing distance from the soma. Notably, we also detected a large variability in [Na^+^]_p_ between different processes of individual cells. Our experiments thus reveal a large intracellular heterogeneity in astrocyte [Na⁺], evident not only between somata and adjacent processes and between different processes of individual cells, but also within an individual process.

### Determinants of baseline [Na^+^] of astrocytes

To analyze likely determinants of astrocytic [Na^+^], we performed dynamic MP-FLIM in bolus-loaded hippocampal slices. Bath application of tetrodotoxin (TTX; 0.5 µM) to block neuronal action potentials neither affected mean somatic [Na^+^]_s_, nor altered its range (control: mean 14.0±6.4 mM, median 13.4 mM, range 6-23 mM; n=53, N=12; TTX: mean 12.7±4.3 mM, median 13.4 mM, range 5-21 mM; n=27, N=9; p=0.6) (Fig. 4a). The potential involvement of gap junctions was studied by bath-application of the gap junction blocker carbenoxolone (CBX; 100 µM). Upon perfusion of CBX, the mean [Na^+^]_s_ of astrocytes as well as its range increased significantly (control: mean 13.0±7.4 mM, median 11.2 mM, range 2.3-45.4 mM; n=186, N=7 CBX: mean 20.0±13.3 mM, median 18.4 mM, range 3.5-96.6 mM; n=134, N=7; p=0.03) (Fig. 4c). As reported before in mice deficient for astroglial Cx43 and Cx30 ^39^, we also found that CBX increased the extracellular K^+^ concentration ([K^+^]_e_) slightly, but significantly from 2.48±0.1 mM to 2.54±0.12 mM as measured using ion-selective microelectrodes at depths of about 50 µm (N=8; p=0.016) (Fig. 4d).

**Fig. 4:**
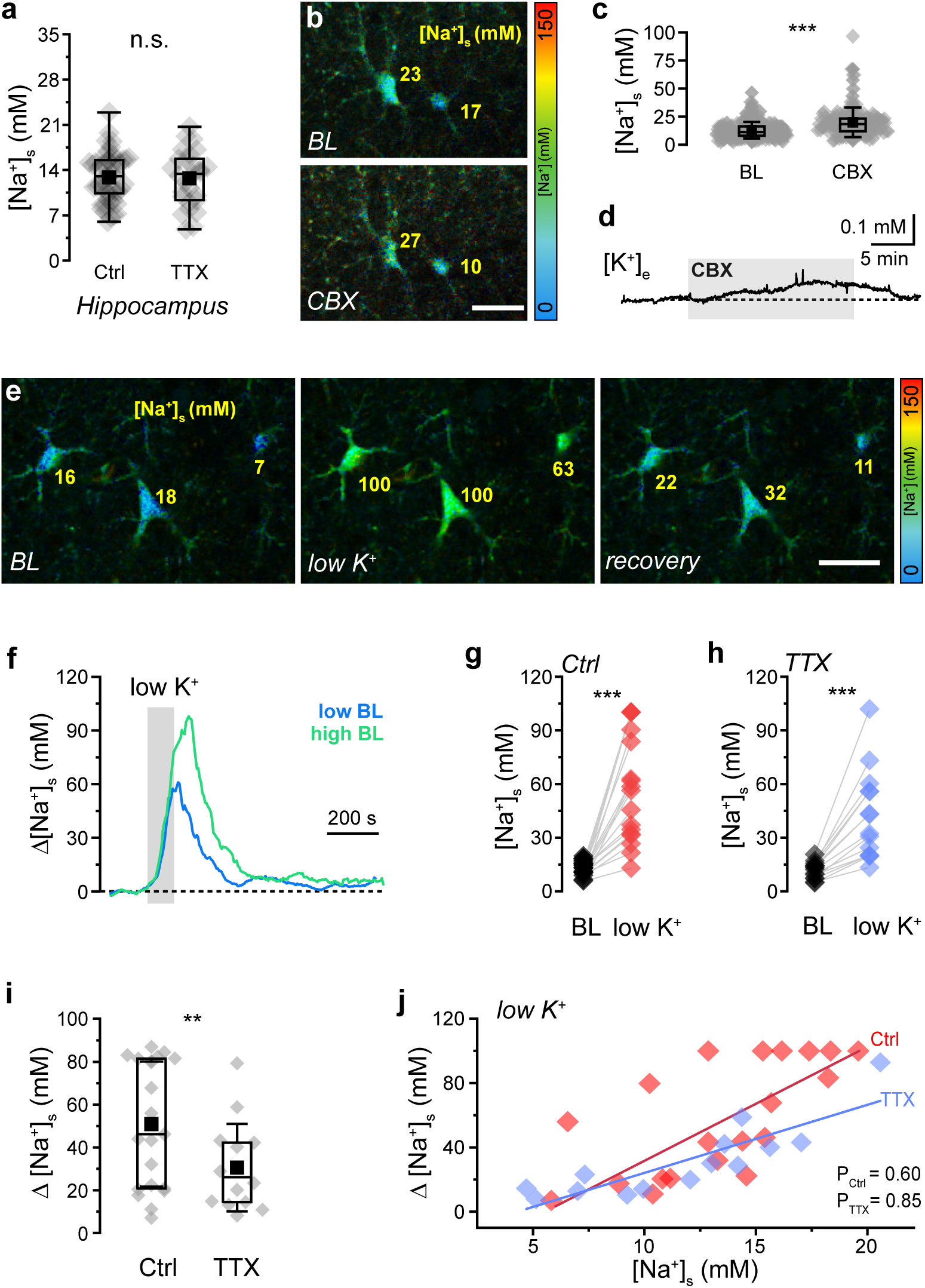
Determinants of astrocytic [Na^+^]. **a** [Na^+^]_s_ under control conditions (Ctrl) and with TTX (Ctrl: n=53, N=4; TTX: n=27, N=5); Mann-Whitney-Test, p=0.60. **b** FL images (masked with SR101) depicting astrocytes at baseline (BL) and after application of carbenoxolone (CBX). **c** [Na^+^]_s_ before (BL) and after addition of CBX (Ctrl: n=186, N=7; CBX: n=134, N=7; Wilcox. Signed Rank test, p=0,03048). **d** Trace illustrating the changes in [K^+^]_e_ induced by CBX (grey area; average of 8 measurements/8 slices). **e-j** Effect of perfusion with K^+^-free saline for 2 minutes (“low K^+^”). **e** FL images (masked with SR101) of an experiment showing baseline [Na^+^]_s_ (BL), peak elevation in [Na^+^]_s_ induced by low K^+^, and [Na^+^]_s_ after recovery. **f** Traces depicting changes in [Na^+^]_s_ induced by low K^+^ in two astrocytes with initially low (7.1 mM, blue trace) and initially high (16.3 mM, green trace) [Na^+^]_s_. **g** [Na^+^]_s_ in astrocytes at baseline (BL) and peak [Na^+^]_s_ induced by low K^+^ (Wilcoxon-signed-rank-test, p=6.4E-5). Lines connect data points from individual cells. **h** same as in g in the presence of TTX (Paired-sample-t test, p=8.82E-5). **i** Peak changes in [Na^+^]_s_ induced by low K^+^ in the absence (Ctrl) and presence of TTX (Mann Whitney Test, p=0.03). **j** Correlation between initial baseline [Na^+^]_s_ and the peak changes in [Na^+^]_s_ induced by low K^+^ without (Ctrl, red) and with TTX (blue) (correlation, Ctrl: Pearson: 0.64, TTX: Pearson: 0.87). **g-j** Ctrl: n=21, N=4; TTX: n=14, N=5. **a, c, g, h, i, j:** diamonds: individual data points, boxes: 25/75, whiskers: SD, lines: median, squares: mean. **b, e:** Color-code for ING-2 FL is shown on the right. Yellow numbers indicate [Na^+^] in the indicated ROIs. All scales: 20 μm.

We next studied the effect of an inhibition of the NKA by perfusing slices with nominally K^+^-free saline for 2 minutes (“low K^+^”) (Fig. 4 e-j). This caused [K^+^]_e_ to transiently decrease to around 1.5 mM (n=8, N=8; not illustrated). At the same time, low K^+^ caused astrocytic [Na^+^]_s_ to rapidly increase to 69.1±31.9 mM (median 73.8 mM, range 13-133 mM; n=24, N=4; p<0.0001) (Fig. 4 e-j). The magnitude of this increase was linearly correlated with the initial baseline [Na^+^]_s_ (Pearson: 0.64, p=0.0018), with astrocytes having a high baseline [Na^+^]_s_ showing a stronger increase than those with a low baseline (Fig. 4 f, g, j). Recovery from low K^+^-induced Na^+^ load followed a monoexponential decay and for cells with an initial baseline [Na^+^]_s_ between 10-15 mM (n=9), the mean decay time constant was 76.6 s. This value increased to 85.7 s for cells with a baseline [Na^+^]_s_ between 15.1-20 mM (n=5), and to 97.4 s for cells with baseline [Na^+^]_s_ between 20.1-31 mM (n=3) (not illustrated). Application of TTX (0.5 µM) reduced the mean peak amplitude of the low K^+^-induced [Na^+^]_s_ increase to 42.5±24.7 mM (median 37.8 mM, range 13-100 mM; n=14, N=5; p=0.02), while maintaining the positive linear correlation between initial baseline [Na^+^]_s_ and increase in [Na^+^]_s_ (Pearson: 0.85, p<0.0001) (Fig. 4 h-j). This data shows that inhibition of the NKA by low [K^+^]_e_ is accompanied by significant Na^+^ influx into astrocytes, which is partly related to action potential firing and neurotransmitter release.

High-affinity Na^+^-dependent uptake of glutamate by EAATs is a major contributor to activity-induced [Na^+^] increases of astrocytes ^14^. To test if EAAT activity influences astrocytic baseline [Na^+^], we perfused hippocampal slices with the EAAT-inhibitor TBOA (1 µM) (Fig. 5 a-d). Perfusion with TBOA caused an elevation in [K^+^]_e_ by about 2.5 mM (n=10, N=10; not shown). Moreover, TBOA resulted in a strong, sustained increase in [Na^+^] in somata of CA1 pyramidal neurons reaching the saturation limit of ING-2 (100-120 mM) within about 6-10 minutes, accompanied by a rounding up and swelling of neuronal cell bodies (Fig. 5 a,b). In contrast, TBOA caused a slow, delayed decrease in astrocytic [Na^+^]_s_ to on average 6.7±3.5 mM (median 5.3 mM; n=36, N=10) with no signs of somatic swelling (Fig. 5 a,b). As observed before, changes in astrocytic [Na^+^]_s_ were weakly linearly correlated with the initial baseline, meaning that astrocytes with a higher baseline exhibited a greater decrease than those with a lower baseline (Fig. 5 c,d).

**Fig. 5:**
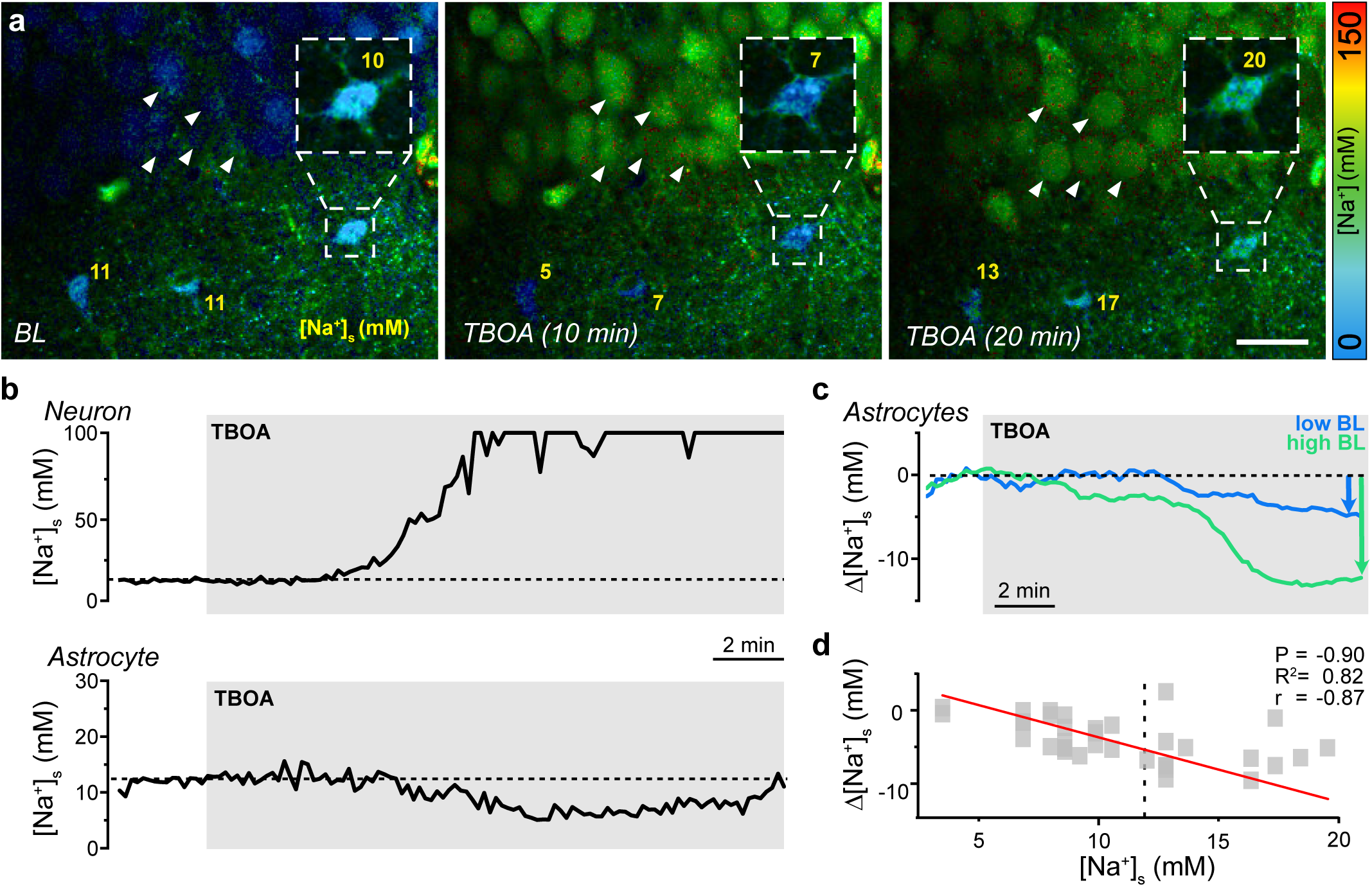
Role of Na^+^-dependent glutamate transporters. **a** FL images of an experiment showing baseline [Na^+^]_s_ (BL), and changes in [Na^+^]_s_ after perfusion with TBOA to inhibit Na^+^-dependent glutamate transporters for 10 and 20 minutes. The box shows the ING-2 FL (masked with SR101) of an astrocyte at higher magnification. Note that cell bodies of CA1 neurons visible in the upper part load with Na^+^ and round up (arrowheads). Color-code for ING-2 FL is shown on the right, yellow numbers indicate [Na^+^] in the indicated ROIs. Scale 20 μm. **b** Traces depicting changes in [Na^+^]_s_ induced by TBOA (grey area) in a CA1 neuron (top) and an astrocyte (bottom). **c** Changes in [Na^+^]_s_ induced in two astrocytes with initially low (12 mM, blue trace) and initially high (20 mM, green trace) [Na^+^]_s_. **d** Correlation between initial baseline astrocytic [Na^+^]_s_ and the peak decrease in [Na^+^]_s_ induced by TBOA (n=36, N=10). Red line: linear fit of the data (Pearson: -0.9, R^2^=0.82).

Taken together, these results show that spontaneous action-potential activity has no detectable influence on baseline [Na^+^]_s_ of hippocampal astrocytes in acute slices. They confirm the expected vital influence of NKA activity on astrocytic [Na^+^]_s_. The data also suggests that cells with a high baseline [Na^+^]_s_ are subject to strong relative influx of Na^+^, whereas cells with a low baseline [Na^+^]_s_ have relatively weaker Na^+^ influx. Alternatively, and/or in addition, high-baseline [Na^+^]_s_ cells might express lower levels of the NKA or display lower NKA activity in response to a Na^+^ load. Moreover, we found that cells with a high baseline [Na^+^]_s_ apparently experience a stronger EAAT-related Na^+^ influx than those with a low [Na^+^]_s_.

### [Na^+^] determines uptake of extracellular K^+^ by astrocytes

Uptake of K^+^ by the astrocytic NKA plays a vital role for homeostasis of [K^+^]_e_ and neuronal excitability ^7^. K^+^-induced activation of the NKA results in well-detectable decreases in astrocyte [Na^+^]_s_ ^40^. To study the influence of [K^+^]_e_ on astrocytic [Na^+^] in the CA1 area of the hippocampus, slices were perfused with ACSF containing 10 mM K^+^ (“high K^+^”) for 2 minutes, resulting in a transient [K^+^]_e_ increase to on average 7.2±0.7 mM (n=13, N=13; not illustrated). This increase in [K^+^]_e_ was approximately three times greater than the increase observed with TBOA (2.5 mM, see above).

In bolus-loaded astrocyte somata, high K^+^ caused an average decline of [Na^+^]_s_ to 6.3±1.6 mM (median: 6.1 mM; n=29, N=8) (Fig. 6 a-f). In individual cells, the magnitude of this decrease showed a strong linear correlation with their former baseline (slope: -0.62; Pearson: - 0.92) (Fig. 6g). For example, cells with a baseline [Na^+^]_s_ of 20 mM showed a decrease more than twice as large as cells with an initial [Na^+^]_s_ of 10 mM (Fig. 6g). The amplitude of high K^+^-induced changes in [K^+^]_e_ was not influenced by TTX (mean 7.1±2.3 mM; n=5, N=5; p=0.53; not illustrated). Moreover, TTX did not alter the high K^+^-induced changes in astrocytic [Na^+^]_s_ (mean 6.6±2.1; n=13, N=4, p=0.2), nor the linear correlation between initial [Na^+^]_s_ and peak decline (slope: -0.53; Pearson: -0.87, p<0.0001) (Fig. 6 d-f). This indicates that the K^+^-induced activation of the astrocytic NKA dominated over any potential Na^+^ influx resulting from increased neuronal activity and/or activation of Na^+^ inward transporters such as EAATs in astrocyte somata.

**Fig. 6:**
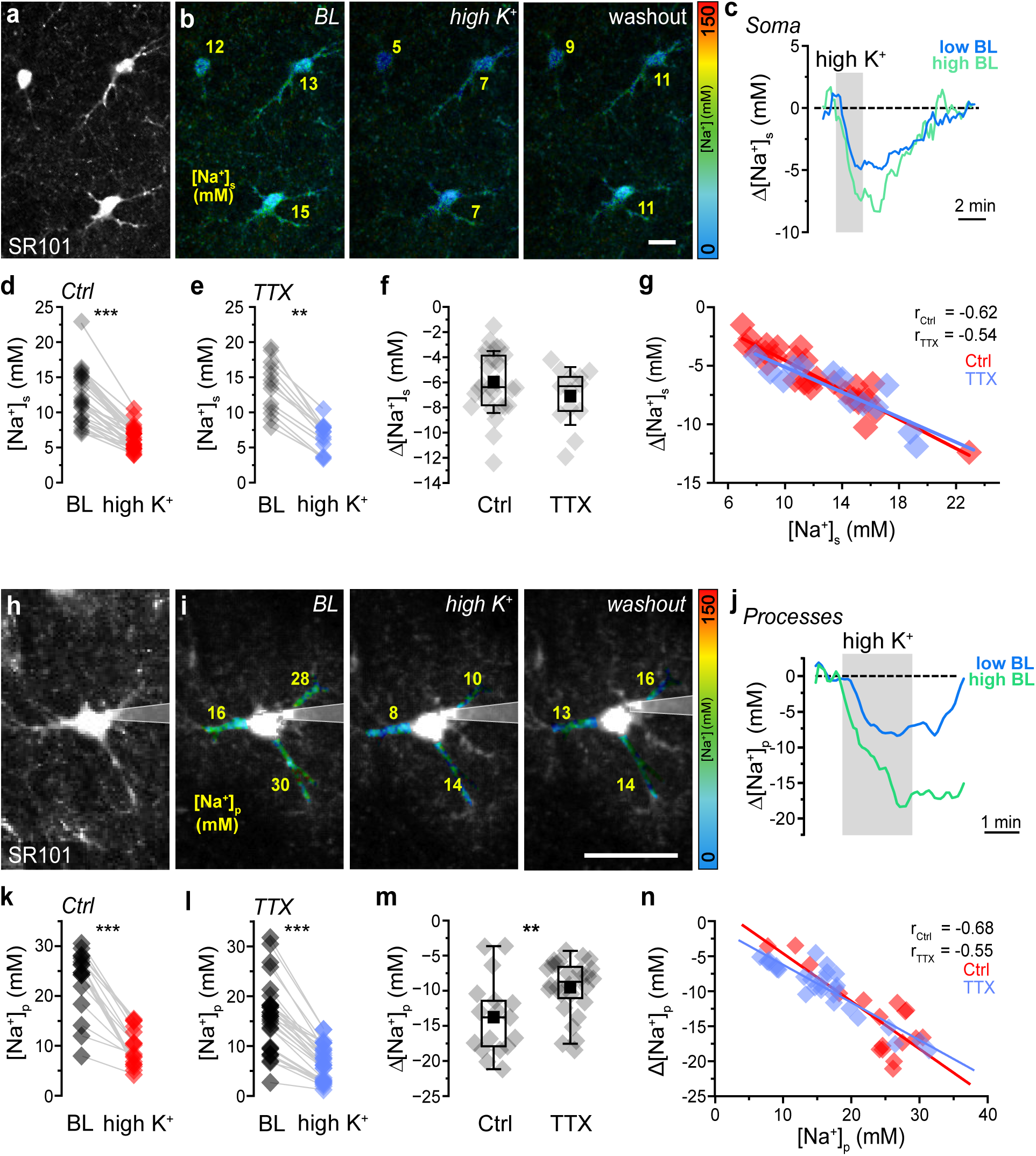
Changes in astrocytic [Na^+^] induced by increasing external K^+^. a-n. Effect of perfusion with saline containing 10 mM K^+^ for 2 minutes (“high K^+^”) on [Na^+^] in somata ([Na^+^]_s_, a-g) and processes ([Na^+^]_p_, h-n). **a** SR101 image depicting several astrocytes. **b** FL images (masked with SR101) of an experiment showing baseline [Na^+^]_s_ (BL), peak decrease in [Na^+^]_s_ induced by high K^+^, and [Na^+^]_s_ after washout of high K^+^ for 5 minutes in three astrocytes. Color-code is shown on the right, yellow numbers indicate [Na^+^] in the indicated ROIs. Scale: 10 μm. **c** Traces showing changes in [Na^+^]_s_ induced by high K^+^ (grey area) in two astrocytes with initially low (8.9 mM, blue trace) and initially high (15.1 mM, green trace) [Na^+^]_s_. **d** [Na^+^]_s_ in astrocytes at baseline (BL) and peak [Na^+^]_s_ induced by high K^+^ (Wilcoxon signed rank test, p=2.70237E-6). Lines connect data points from individual cells. **e** same as in d but with TTX present (Wilcoxon signed rank test, p=0.00166). **f** Peak decrease [Na^+^]_s_ induced by high K^+^ in the absence (Ctrl) and presence of TTX (Mann-Whitney Test, p=0.21). **g** Correlation between initial baseline [Na^+^]_s_ and the peak decrease in [Na^+^]_s_ induced by high K^+^ without (Ctrl, red) and with TTX (blue). Lines represent linear fits of the data (correlation, Ctrl: Pearson=-0.92; TTX: Pearson=-0.88). **a-g:** Ctrl: n=29, N=8; TTX: n=13, N=4. **h-n** Same as in a-g, but for processes ([Na^+^]_p_) from astrocytes dye-loaded by whole-cell patch-clamp. **i** Time series (binning 5) of summed extended focus FL-images (3 z-steps, 3.5 µm step size). Right image shows [Na^+^]_s_ after washout of high K^+^ for 2 minutes. **j** Traces from two processes with initially low (16 mM, blue) and high (28 mM, green) baseline [Na^+^]_p_. **k-l** Paired-sample-t test, Ctrl: p=1.53E-05; TTX: p=1.28E-12. **m** Mann-Whitney-Test: p=0.00861. **n** Correlation, Ctrl: Pearson: -0.90; TTX: Pearson: -0.82. **h-n:** Ctrl: n=17, N=5; TTX: n=27, N=5. **d-g, k-n:** diamonds: individual data points, boxes: 25/75, whiskers: SD, lines: median, squares: mean.

To analyze the influence of high K^+^ on [Na^+^] in individual processes, astrocytes were dye-filled by whole-cell patch-clamp (Fig. 6 h,i). Increasing [K^+^]_e_ caused an average decrease of [Na^+^]_p_ from 23.3±6.5 mM to 9.5±3.7 mM (Fig. 6 j,k), which is significantly larger as compared to the soma (n=17, N=5; p<0.0001). During perfusion of TTX, high K^+^ reduced average [Na^+^]_p_ from 16.6 ± 6.5 mM to 7.0 ± 3.4 mM (n=27, N=5; p=0.009) (Fig. 6l). As opposed to somata, TTX thus decreased the mean amplitude of the high K^+^-induced decline in [Na^+^]_p_ (p=0.009, Fig. 6m), suggesting that high K^+^ induced action-potential-related neuronal activity promoted the drop in astrocyte [Na^+^]_p_. As observed for somata, average [Na^+^]_p_ and the high K^+^-induced decline in [Na^+^]_p_ followed a linear correlation with a comparable slope in the presence and absence of TTX (control: slope -0.68, Pearson: -0.81, p<0.0001; TTX: slope -0.53, Pearson: -0.87, p= 0.00013) (Fig. 6n).

Taken together, these results show that an increase in [K^+^]_e_ results in an immediate decrease in [Na^+^] in somata and processes of astrocytes, the magnitude of which is strongly correlated with the initial baseline [Na^+^]. As Na^+^ export is virtually solely mediated by the NKA, this suggests that compartments with a high baseline [Na^+^] undergo a stronger K^+^-induced activation of the NKA than those with a low [Na^+^].

### Biophysical modeling of astrocytic [Na^+^]

Astrocytes express different NKA subunits, and recent RNA sequencing showed predominant expression of α2β2 in about 70% of mouse forebrain astrocytes, while about 30% express α2β1 ^35,41,42^. As these differ in their binding affinities for internal Na^+^ and external K^+^ ^6^, we performed mathematical simulations to study the influence of a specific subunit composition on astrocytic baseline [Na^+^] and changes in [Na^+^] induced by low or high K^+^.

For a fixed NKA expression level and Na^+^ influx, these simulations predict a baseline [Na^+^] of 11.8 mM for cells with α2β1 subunits only (Fig. 7a, simulation 1). Exclusive expression of α2β2 resulted in a [Na^+^] of 22.3 mM, whereas mixing both NKA isoforms resulted in levels in between (Fig. 7a, simulation 1). Next, we fixed the isoforms to the reported distribution of 30% α2β1/70% α2β2 ^41,42^, and varied the overall NKA expression level, i.e. pump strength, between 60-180%. Baseline [Na^+^] increased with decreasing NKA expression from ∼8 mM at 180% to >40 mM at 60% (Fig. 7a, simulation 2). Varying the strength of Na^+^ influx between 50-170% at fixed NKA composition (30% α2β1/70% α2β2) and expression level, caused baseline [Na^+^] to increase from ∼8 mM to near 40 mM (Fig. 7a, simulation 3). Finally, we performed simulations to obtain a best fit of the experimentally determined bimodal distribution of [Na^+^] (see Fig. 1h), varying the NKA isoform composition (α2β1/α2β2) and pump strength (80-180%). We found that a combination of these parameters can replicate the range and distribution profile of astrocyte [Na^+^]_s_ reported by the FLIM measurements (Fig. 7a, simulation 4).

**Fig. 7:**
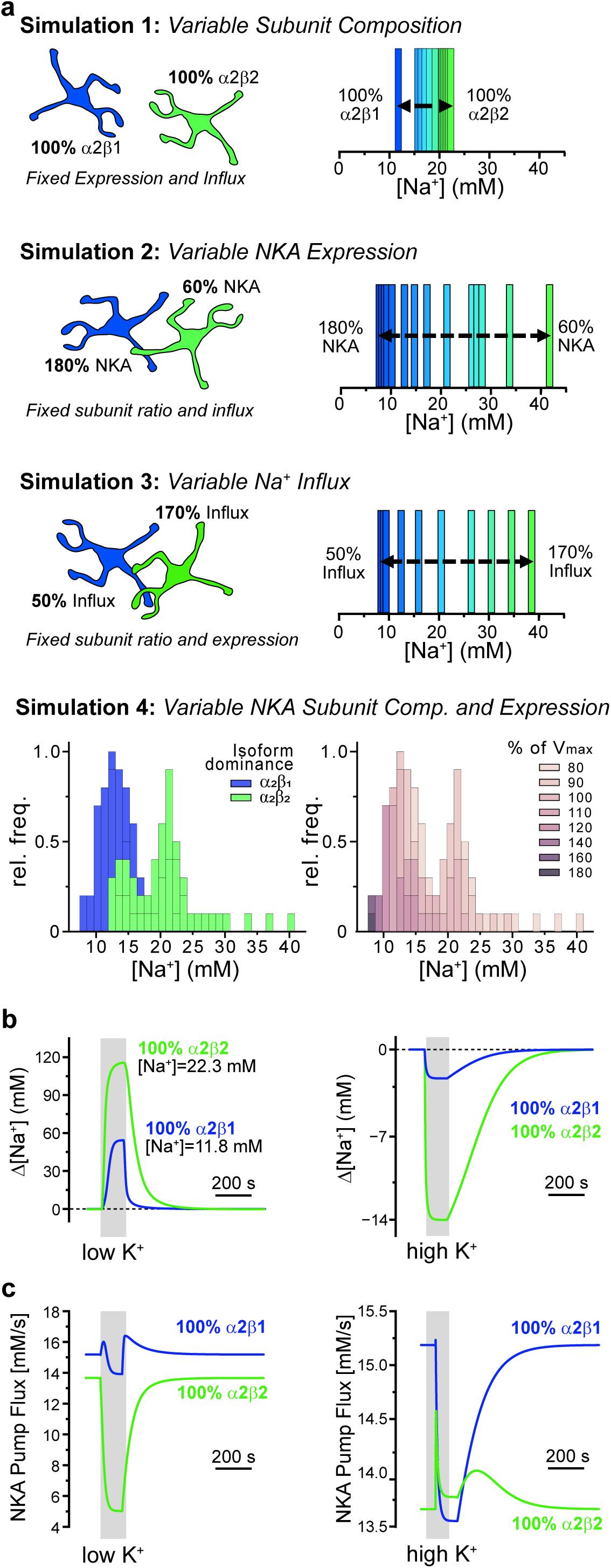
Biophysical modeling of astrocytic [Na^+^]. **a** Results from simulations illustrating the determinants of [Na^+^]. Simulation 1: Influence of varying the NKA subunit composition between 100% α2β1 to 100% α2β2 on astrocytic baseline [Na^+^] while keeping NKA expression levels and Na^+^ influx constant. Simulation 2: Variation of NKA expression levels/pump strength from 180% to 60% at fixed Na^+^ influx and NKA subunit composition (α2β1/α2β2 = 30/70). Simulation 3: Variation of Na^+^ influx from 50% to 170% at fixed NKA subunit composition (α2β1/α2β2 = 30/70) and fixed NKA expression levels/pump strength (100%). Simulation 4: Simulation combining variable isoform dominance with variable NKA expression level/pump strength as indicated. Left: Bar-chart illustrating whether α2β1 (blue) or α2β2 (green) expression dominates. Right: Same bar chart as left, but color-code indicating relative NKA expression level/pump strength (V_max_). **b** Left: Simulation showing the changes in [Na^+^] induced by K^+^-free saline for 2 minutes (“low K^+^”) in astrocytes with exclusive expression of α2β1 (blue) or α2β2 (green) NKA isoforms. Right: simulation showing changes in [Na^+^] induced by an increase in [K^+^]_e_ to 10 mM (“high K^+^”). **c** Time traces of flux through the two isoforms of NKA during the low [K^+^]_e_ simulation (α2β1: blue trace, α2β2: green trace) with low K^+^ (left) or high K^+^ (right).

Next, we tested how a specific NKA composition influences the astrocytes’ response to NKA inhibition by subjecting them to 1 mM [K^+^]_e_ for 2 minutes. Astrocytes with exclusive α2β1 expression showed a [Na^+^] increase by ∼55 mM, while cells with α2β2 experienced an increase by >100 mM (Fig. 7b, left). In resting state, the flux through α2β1 is higher than α2β2, resulting in lower intracellular [Na^+^] (Fig. 7c, left; see also Fig. 7a, simulation1). Under low K^+^ condition, the flux through α2β1 first increases as intracellular [Na^+^] rises, followed by a smaller decrease due to the drop in [K^+^]_e_ as compared to α2β2 (Fig. 7c, left). Overall, this led to a smaller increase in intracellular [Na^+^] in α2β1 astrocytes as compared to astrocytes expressing α2β2 (Fig. 7b, left). Finally, we analyzed the relevance of subunit compositions for activation of the NKA by external K^+^. Here, an opposite behavior can be seen, where the flux through α2β2 increases as we increase [K^+^]_e_ causing the intracellular Na^+^ to drop more (Fig. 7 b,c, right). α2β1 on the other hand is already close to saturation and its flux increases only slightly as [K^+^]_e_ rises. However, it decreases sharply in response to the drop in intracellular [Na^+^], countering a further decrease in [Na^+^] (Fig. 7 b,c, right). Overall, this leads to a smaller decrease in [Na^+^] in α2β1-expressing astrocytes (Fig. 7b, right). In Figure S1, and the interactive 3D-Plots (<u>Link</u>), we show how the flux through the two isoforms changes as we vary [K^+^]_e_ and [Na^+^] further.

Taken together, these simulations demonstrate that the specific subunit composition of the NKA, together with its expression level and the magnitude Na^+^ influx are important determinants of the baseline [Na^+^] of astrocytes. Moreover, they show that a combination of these parameters can reproduce the relatively broad, biphasic distribution of astrocyte [Na^+^] determined experimentally. In addition, the simulations replicate the experimentally observed dependence of the magnitude of [Na^+^] changes induced by inhibition of the NKA as well as its stimulation by extracellular K^+^.

## Discussion

Employing MP-FLIM, we performed the first quantitative analysis of the intracellular [Na^+^] of astrocytes in mouse forebrain tissue slices and in the mouse cortex *in vivo* ^33^. Our measurements revealed a mean somatic [Na^+^] of 13-14 mM in astrocytes *in situ*, which is similar to values reported using intensity-based imaging (e.g., ^43–48^). Somatic [Na^+^] of layer 2/3 cortical astrocytes *in vivo* was around 20 mM and thus significantly higher, a difference likely related to higher levels of neuronal activity (e.g., neurotransmitter release and uptake) in the intact brain compared to slices.

Both *in situ* and *in vivo*, somatic [Na^+^] exhibited a relatively broad range. Na^+^ is a highly mobile, unbuffered ion, suggesting its equilibration within astrocytes and between gap-junction-coupled cells at rest ^18,20,49^. Inhibition of gap junctions with carbenoxolone increased the overall range of [Na^+^]_s_ in hippocampal astrocytes, indicating a diffusion-driven exchange of Na^+^ between cells as reported from astrocytes in culture ^49^. Moreover, carbenoxolone caused an increase in the average [Na^+^]_s_, which could be related to increased neuronal excitability, a phenomenon described in mice deficient for astroglial Cx30 and Cx43 ^50^. This idea is in line with the slight increase in [K^+^]_e_ induced by carbenoxolone, also reported from astrocyte-specific Cx43/Cx30-deficient mice ^39^. However, unspecific effects cannot be ruled out as carbenoxolone has also direct effects on neurons ^51,52^.

FL-based imaging of a large number of cells revealed that in hippocampal slices, astrocytic [Na^+^]_s_ follows a bimodal distribution with two Gaussian components that peak at 8 and 16 mM. A similar biphasic distribution has been described for basal [Ca^2+^] of hippocampal astrocytes *in situ* and of cortical astrocytes *in vivo* ^31^. These observations suggest the existence of at least two distinct functional subgroups, supported by the reported molecular heterogeneity of astrocytes ^35,37,38,53–55^. Notably, this molecular heterogeneity includes NKA isoforms. Rodent forebrain astrocytes predominantly express the α2 isoform, which is combined with β1 in approximately 10–30% of cells, while the remainder expresses β2 ^35,56^. Based on their different ion binding affinities ^7,56^, these two NKA subunit compositions are associated with differences in baseline [Na^+^] as also demonstrated by our simulations.

The experimentally observed heterogeneity in astrocytic [Na⁺] is thus likely to be due, at least in part, to different NKA isoforms, with exclusive α2β1 expression resulting in low (∼12 mM) and α2β2 in high baseline [Na^+^] (∼22 mM). Altering the overall expression/maximum NKA pump strength in our simulations from 60-180% caused this range to increase even further (from ∼8 to >40 mM), indicating that different NKA expression levels also contribute to the broad distribution of basal [Na^+^]. Finally, both experiments and simulations strongly suggest that differences in Na^+^ influx rates influence [Na^+^]. Following NKA inhibition, cells with a high baseline [Na^+^] (α2β2) exhibited a larger increase in [Na^+^], than those with a low baseline [Na^+^] (α2β1), indicating stronger Na^+^ influx combined with overall lower NKA activity in the former.

Our experiments also revealed a substantial subcellular heterogeneity in [Na^+^]. [Na^+^]_p_ not only varied between different main processes of a given cell, but also increased with increasing distance from the soma, a pattern comparable to the radial gradient described for [Ca^2+^] in astrocyte processes ^31^. These findings are surprising due to the high ionic mobility of Na^+^ and the absence of Na^+^-buffer systems ^14^, but in line with modeling studies suggesting the existence of intracellular domains and subcellular gradients of Na^+^, caused by the restricted flow and retention of cations in fine astrocyte processes ^22,23^. Increased [Na^+^]_p_ is likely to result from increased Na^+^ influx following local neuronal activity and release of transmitters, shown to locally depolarize astrocytic processes ^24^. Our experiments suggest that EAATs are a major mediator of Na^+^ influx as their inhibition with TBOA caused astrocytic [Na^+^]_s_ to decline. The magnitude of this decline depended on the initial [Na^+^], as high [Na^+^]-astrocytes showed a greater fall in [Na^+^] than those with a low initial [Na^+^]. In this context, it is noteworthy that the increase in [K^+^]_e_ observed with TBOA was only a fraction of that seen with high K^+^ perfusion, whereas the TBOA-induced decrease in [Na^+^] was greater, suggesting that the observed effects were indeed predominantly attributable to decreased Na^+^ influx following EAAT inhibition. This conclusion is in line with former work identifying EAATs as the main drivers of intracellular Na^+^ transients induced by synaptic stimulation in hippocampal astrocytes ^19,45^.

EAATs are predominantly expressed in astrocyte processes and perisynaptic regions and their surface density increases from the soma towards the tips of processes ^57–61^. The observed heterogeneity in [Na^+^]_p_ is thus likely due to the spatial heterogeneity in EAAT expression. In other words, the observed increase in [Na^+^]_p_ from the soma towards the periphery may be generated, at least in part, by increased EAAT-mediated Na⁺ entry into distal versus proximal processes. In addition, the variable proximity of astrocyte processes to glutamate-releasing synapses will contribute to the heterogeneity in [Na^+^]_p_ ^26,27^. EAATs are co-localized with α2-NKA in astrocyte processes, which is also most highly expressed close to synapses ^12,57,58,62^. Intriguingly, the intracellular Na^+^ gradient along processes as demonstrated here will enable the directed diffusion of Na⁺ from the periphery towards the large-volume soma, thereby promoting recovery from activity-related Na^+^ increases in perisynaptic astrocytic processes. Of note, such fast, diffusion-driven recovery will reduce the local ATP consumption through NKA-related export of Na^+^ ^18,19,63^.

We also found that baseline [Na^+^] of a given compartment determined its response to increases in bath [K^+^]_e_, which are mainly cleared by K^+^-induced activation of the NKA ^7,8^. High-baseline [Na^+^] astrocytes showed a stronger decline in [Na^+^] upon [K^+^]_e_ elevation than low-baseline [Na^+^] astrocytes, suggesting that compartments with a high baseline [Na^+^] undergo a stronger K^+^-induced activation of the NKA than those with a low [Na^+^]. Due to its lower affinity for extracellular K^+^ (K_m_ ∼3.6 mM), the α2ß2 combination is specifically geared towards uptake of K^+^. In contrast, α2ß1 has lower affinity for intracellular Na^+^ (K_m_ ∼10.6 mM) and is thus more efficiently activated by [Na^+^] increases ^6,7,56^. Indeed, our simulations demonstrate that the NKA-related pump current increased in α2β1 astrocytes in response to Na^+^ loading, indicating its stimulation by increased [Na^+^]; this was not the case in α2β2 cells. Moreover, α2ß1 astrocytes showed a much smaller decrease in [Na^+^] in response to increases in bath [K^+^]_e_ than α2ß2 astrocytes. This is in line with our experimental results which demonstrated that the magnitude of the K^+^-induced [Na^+^] decrease, and NKA activation, respectively, was positively correlated with the initial baseline [Na^+^]. This again clearly indicates that low baseline [Na^+^] astrocytes indeed predominately express α2ß1, whereas high-baseline [Na^+^] astrocytes predominately express α2ß1.

In conclusion, our study reveals a hitherto unexpected cellular and subcellular heterogeneity in astrocyte baseline [Na^+^]. We demonstrate that the broad range of astrocytic [Na^+^] can be explained by the combination of specific NKA isoforms (α2ß2 versus α2ß1) and expression levels, as well as by differences in the Na^+^ influx via transporters such as EAATs. Notably, these players also determine the strength of NKA activation and of uptake of K^+^ uptake and glutamate from the extracellular space. Astrocytes which display a high [Na^+^], indicative of expression of α2ß2 and a high rate of Na^+^ influx via EAATs and other Na^+^-dependent transporters, are thus especially geared towards an efficient clearance and control of both [K^+^]_e_ and extracellular glutamate, shaping and controlling the excitability of surrounding neurons and networks.

## Methods

### Animals

The study was conducted in accordance with the guidelines of the Heinrich Heine University Düsseldorf (HHU), as well as the European Community Council Directive (2010/63/EU) for *in situ*, and Directive (86/609/EEC) for *in vivo* measurements, as well as all relevant national and institutional guidelines and requirements. Mice were housed under 12 h light/dark conditions with food and water access *ad libitum*. All experiments using brain tissue slices were communicated and approved by the Animal Welfare Office at the Animal Care and Use Facility of the HHU (Institutional Act No. O52/05). For *in vivo* experiments, all procedures were approved by the Landesamt für Natur, Umwelt und Verbraucherschutz Nordrhein-Westfalen (LANUV, Germany), where required.

### Experimental design and statistical analysis

Each set of experiments was performed on at least three different animals. *In* situ experiments were performed on at least 4 different slices obtained from mice of both sexes. *In vivo* imaging was performed on 3 male mice. *n* represents the number of analysed cells, *N* represents the number of slice preparations *in situ*.

The data was tested for normal distribution using the Shapiro-Wilk test. Normally distributed paired data was statistically analysed by a paired t-test. Unpaired parametric data was analysed with a t-test when comparing two groups, and with a one-sided ANOVA with Bonferoni post-hoc correction if more than two groups were tested. Not normally distributed paired data was analysed with a Wilcoxon signed rank test, and unpaired data with a Mann-Whitney-Test for significance. For a detailed list of which tests were used for which data set and the statistical outcome see the Supplemental Statistical Summary. The results of the statistical tests are illustrated as follows: *: 0.01 ≤ p < 0.05; **: 0.001 ≤ p <0.01 and ***: p < 0.001.

### Preparation, salines and drug application

Brain tissue slices were prepared from BALB/c mice (both sexes) as described before (for details see ^64^). In brief, mice of postnatal days (P) 14–20 were anesthetized with CO_2_ based on the recommendations of the European Commission ^65^, rapidly decapitated and their brains removed. Hemispheres were separated and cut into 250-µm-thick slices, while placed in ice-cooled preparation saline (pACSF), containing (in mM): 130 NaCl, 2.5 KCl, 0.5 CaCl_2_, 6 MgCl_2_, 1.25 NaH_2_PO_4_, 26 NaHCO_3_, and 5 glucose, bubbled with 95% O_2_/5% CO_2_, pH 7.4, and an osmolarity of 310 mOsm l^-1^. After cutting, slices were incubated at 34°C for 20 min in pACSF, containing 0.5-1 µM sulforhodamine 101 (SR101) to selectively stain astrocytes. Subsequently, they were transferred to standard artificial cerebrospinal fluid (ACSF), containing (in mM): 130 NaCl, 2.5 KCl, 2 CaCl_2_, 1 MgCl_2_, 1.25 NaH_2_PO_4_, 26 NaHCO_3_ and 5 glucose, bubbled with 95% O_2_/5% CO_2_, pH 7.4, and an osmolarity of 310 mOsm l^-1^ for 10 minutes at 34°C, after which slices were kept in ACSF at room temperature (21±1° C) for up to 6 hours.

During experiments, slices were continuously perfused with ACSF at room temperature. To alter [K^+^]_e_, slices were perfused with ACSF, in which the [K^+^] was adjusted to 0 mM (“low K^+^”) or 10 mM (“high K^+^”), while keeping [Na^+^] + [K^+^] at 160 mM. Pharmacological blockers (tetrodotoxin (TTX, 0.5 µM, HLB-HB1035, Biozol, Hamburg, Germany); carbenoxolone (CBX, 100 µM, C4790-5G, Sigma Aldrich); DL-threo-β-Benzyloxyaspartic acid (TBOA, 1 µM, CAS: 205309-81-5, Tocris, Bio-Techne, MN, USA) were added to the ACSF and applied by bath perfusion.

For calibration of ING-2 FL, slices were perfused with calibration salines containing (in mM): 10 HEPES, 26 K-gluconate, 0-150 Na^+^, 0-150 K^+^ (total concentration of NaCl + KCl was 150 mM), pH was adjusted to 7.4 with KOH. In addition, calibration salines contained 3 µM gramicidin, 10 µM monensin and 100 µM ouabain (Calbiochem, Merck KGaA Darmstadt, Germany) to permeabilize cellular plasma membranes for Na^+^ as described earlier ^45,66^.

### Fluorescence lifetime imaging *in situ*

For Na^+^ imaging, the membrane-permeable form of the chemical Na^+^ indicator ION-NaTRIUM-Green-2-AM (ING-2; Mobitec GmbH, Göttingen, Germany, #2011F) was pressure-injected as reported before ^33^. Multi-photon fluorescence lifetime imaging microscopy (MP-FLIM) of ING-2 was performed using a modified laser-scanning microscope based on a Nikon A1-R MP system (Nikon Europe, Amsterdam, The Netherlands), equipped with a water immersion objective (NIR Apo 60x/NA 1.0, Nikon). Laser pulses (<100 fs, 840 nm) were generated at 80 MHz by a mode-locked Titan Sapphire laser (Mai Tai DeepSee, Newport, Spectra Physics; Irvine, CA, USA).

Images were acquired at ∼1 Hz and temporally binned depending on the needed temporal resolution. Average fluorescence lifetimes (FL) were measured using time-correlated single photon counting (TCSPC) with a spatial resolution of 0.41x0.41 µm per pixel. Fluorescence emission was split with a 560 nm long pass dichroic mirror (H 560 LPXR, F48-562 AHF Analysentechnik AG, Tübingen, Germany) and band-pass filtered at 540/25 nm (F34-540A, AHF) for ING-2 and at 640/20 nm (F39-641, AHF) for SR101, before being directed to PMA hybrid photodetectors (PicoQuant, Berlin, Germany). TCSPC electronics (Multiharp 150, PicoQuant) and acquisition software (Symphotime64, Version 2.6, PicoQuant) were used for obtaining FL at a pixel dwell time of 3.81 µs for frames consisting of 512x512 pixels. Astrocytes were selected in the SR101 image and only those with a total photon count of >5000 photons and at least 5 photons per pixel per frame were used for analysis. Acquired images were analysed using Symphotime 64 software (Version 2.9, PicoQuant). To dynamically image astrocytic processes, XYZT stacks of variable size (3-5 steps) were taken at a temporal binning of 5 frames for each step. The XYZT stack was summed in Z-axes prior to the analysis of individual processes to yield a sufficient number of photons for lifetime calculation.

For calibration of ING-2 FL, the relationship between [Na^+^] and amplitude-weighted average decay constant τ = (A1 * τ1+A2 * τ2)/(A1+A2) was approximated by a shifted Michaelis-Menten kinetics function τ = (τ_Max_ * [Na^+^])/(K_m_+[Na^+^])+τ_min_, where τ_min_ corresponds to [Na^+^]=0 mM, and τ_Max_ as well as K_m_ are computed from the fitted function using OriginPro 2024b (OriginLab Corporation, Northampton, USA). Average FL data from *in situ* experiments was constrained to the boundaries of the calibration and dynamic range of the dye, respectively (lower limit: 0 mM and upper limit: 100 mM Na^+^, compare Fig. 1e).

### *In vivo* MP-FLIM and analysis

*In vivo* MP-FLIM was performed in layer 2/3 of the barrel cortex in postnatal week 8-12 C57BL/6 male mice, as previously described ^32,67,68^. Animals were anesthetized using isoflurane inhalation (3.5% for induction, 1.2–1.5% for maintenance) and additional buprenorphine analgesia (0.1 mg/kg). The mouse’s head was fixed in a stereotaxic apparatus, the skull exposed, and a small craniotomy (3 mm diameter) above the right barrel cortex was performed. The skull was carefully removed and the dura matter kept intact. Then, 300 nl of a solution containing 750 µM ING-2 AM, Pluronic-127F (10% in DMSO) and sulforhodamine 101 (SR101, 25 µM) dissolved in artificial cerebrospinal fluid (152 mM NaCl, 2.5 mM KCl, 2 mM CaCl_2_, 10 mM HEPES, 1.25 mM NaH_2_PO_4_, 1 mM MgSO_4_, 10 mM D-glucose, pH 7.4) was pressure-injected via a glass micropipette at a depth of 120-150 µm from the dura mater. The eyes were covered with Bepanthen (Bayer Vital GmbH, Leverkusen, Germany) and animals were kept at a constant temperature of 37°C throughout surgery and imaging experiments.

Recordings were performed under continued isoflurane inhalation anesthesia (1.0–1.2%), for which mice were kept head-fixed and placed under a two-photon excitation fluorescence microscope (COSYS Ltd, East Sussex, UK) equipped with a femtosecond infrared pulsed Mai Tai HP laser (Spectra-Physics) at 840 nm and a 16× water-immersion objective lens (Nikon LWD, NA 0.8). Images (512×512 pixels) were acquired at a depth of 100–150 μm from the pia with a nominal resolution of ∼0.4 μm/pixel. Emitted light was spectrally separated using an appropriate dichroic mirror and filter set (see methods section for *in situ*).

Astrocytes were identified using SR101 and imaged in intensity mode using ScanImage 2022.1.0 (MBF Bioscience, Williston, USA). ING-2 AM was imaged in TCSPC mode using MultiHarp 150 module (PicoQuant) and data were acquired with SymPhoTime 64 software (PicoQuant). Frame scans (20-100 frames) were acquired in galvo-galvo scanning mode at a frame rate of 0.95 Hz. Prior to analysis, *in vivo* data were motion-corrected where appropriate and analyzed offline using in-lab-written MATLAB (The MathWorks, Natick, USA) scripts. Calibration of ING-2 lifetimes was performed in a cuvette using calibration solutions as described above, but free of ionophores and ouabain. Minor differences between fluorescence lifetime components observed in the cuvette and *in vivo* were accounted for as previously described ^32^.

### Image processing for illustrations

All images presenting τ_AVG_ are fast lifetime images and were processed equally for illustration purposes regarding adjustment of brightness and contrast within each respective figure. To better visualize astrocyte processes, the fast lifetime images were masked with SR101 fluorescence intensity as shown in Figure S2 and as indicated in the respective figure legends. This effectively masks areas that appear dark in the SR101 intensity image, thereby increasing the visual perception of astrocyte processes.

### Electrophysiology

Individual astrocytes were subjected to whole-cell patch clamp using an EPC10 amplifier and “PatchMaster NEXT” software (MCS GmbH/HEKA Elektronik, Reutlingen, Germany). Patch pipettes with a resistance of 2.0-3.5 MΩ were pulled from borosilicate glass capillaries (GB150(F) 8P, Science Products, Hofheim am Taunus, Germany) using a vertical puller (PC-10 Puller, Narishige International, London, UK). The pipette solution contained (in mM): 116 K-Gluconate, 32 KCl, 10 HEPES (N-(2-hydroxyethyl)piperazine-N′-2-ethanesulfonic acid), 10 NaCl, 4 Mg-ATP, 0.4 Na_3_-GTP and 0.05 ING-2 (MoBiTec; Göttingen, Germany), the pH was adjusted to 7.3, the osmolarity was 305 ± 5 mOsm. Cells were held in the whole-cell mode for a minimum of 15 minutes prior to starting imaging experiments. Cells were excluded from analysis if the input resistance exceeded 25 MΩ. Data was analyzed using “OriginPro 2024b” (OriginLab Corporation, Northampton, MA, USA).

Double-barreled ion-sensitive microelectrodes were employed to measure [K^+^]_e_ as described before ^69^. In brief, two borosilicate glass capillaries with filament (GC100F-15, GC150F-15; Harvard Apparatus, Holliston, MA, USA) were glued and pulled out together. The tip of the K^+^-sensitive barrel was filled with valinomycin (Ionophore I, Cocktail B, Merck, Darmstadt, Germany) and backfilled with 100 mM KCl. Reference channels were filled with HEPES-buffered saline containing (in mM): 125 NaCl, 3 KCl, 25 HEPES, 2 MgSO_4_, 2 CaCl_2_, 1.25 NaH_2_PO_4_ and 10 glucose; pH 7.4. Calibration of K^+^-sensitive microelectrodes was performed in the bath using salines composed of 25 mM HEPES and a total of 150 mM NaCl and KCl, in which [K^+^] was 0–10 mM and [Na^+^] adjusted accordingly. After calibration, electrodes were positioned in the *stratum radiatum* at ∼50 µm below the slice surface and measurements performed. Electrodes were calibrated again in the experimental bath directly after each experiment.

### Simulations

The equations and related parameters modelling the dynamics of membrane potential and ion concentrations in the astrocyte and extracellular space are described in the Supplementary Information. The rate equations were integrated using Euler method in Python 3.

## Data Availability

The datasets generated during and/or analyzed during the current study are available from the corresponding author on reasonable request.

## Code availability

The code used to compute the results and statistics, as well as generate visualizations is deposited in a GitHub public repository at https://github.com/banalok/Modeling

## Acknowledgements

This study was supported by the Deutsche Forschungsgemeinschaft (DFG, German Research Foundation, Projekt #461542557 to C.R.R. and HE6949/8 to C.H.), the Federal Ministry of Education and Research (BMBF), Germany (Project SynGluCross to C.R.R. and C.H.) and by the National Institutes of Health (R01NS130916 and R21AG087910 to G.U.). The authors wish to thank Claudia Roderigo and Simone Durry for expert technical assistance.

## Author contributions

Conceptualization, C.R.R., J.M. and G.U..; methodology, J.M., S.E., P.U., C. H., K.W.K., S.D., and V.B.; experiments, formal and biostatistical analysis: J.M., S.E., P.U., C.H. and V.B; biophysical modeling, A.B. and G.U.; writing – original draft, review and editing, all authors; supervision: C.R.R., C.H. and G.U.; funding acquisition, C.R.R., C.H. and G.U..

## Materials & Correspondence

Correspondence and requests for material should be addressed to Christine R. Rose (slice experiments), Christian Henneberger (*in vivo* experiments) or Ghanim Ullah (modeling).

---

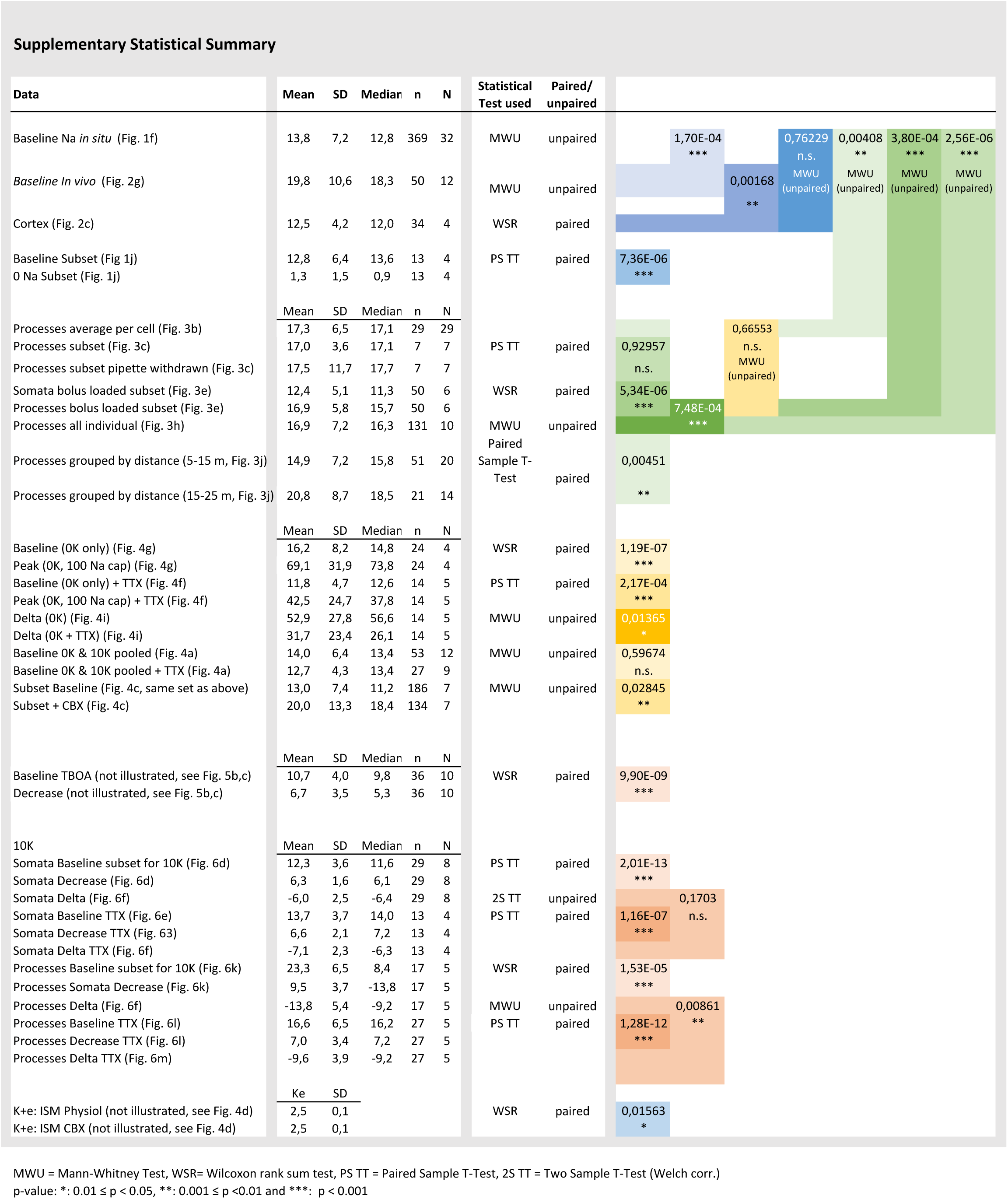

## Supplementary Information

### Model equations for the dynamics of membrane potential and various ion concentrations in the astrocytes and extracellular space

The astrocyte model is adapted from [1,2,3,4] and focuses on the dynamics of various ionic channels, transporters, and cotransporters involved in astrocytic function. The Na⁺/K⁺-ATPase (NKA) maintains ionic homeostasis by actively exporting three Na⁺ ions in exchange for two K⁺ ions at the cost of ATP. In astrocytes, it works alongside other critical transporters to regulate ion dynamics and pH. The electrogenic Na^+^-HCO_3_^-^ cotransporter 1 (NBCe1) contributes by co-transporting Na⁺ and HCO_3_^-^ into the cell, aiding in intracellular pH regulation and influencing Na⁺ accumulation. The Na⁺/H⁺-exchanger (NHE) further modulates pH and Na⁺ levels by importing Na⁺ while exporting protons. The model also includes glutamate transporters that mediate the uptake of glutamate into astrocytes, facilitating the recycling of

neurotransmitters and preventing excitotoxicity. To regulate intracellular Ca²⁺, Na⁺/Ca^2+^-exchangers (NCX) and Transient Receptor Potential Vanilloid 4 channels (TRPV4) are included. In forward mode, NCX removes excess Ca²⁺ while importing Na⁺, and TRPV4 contributes to Ca²⁺ influx.

The model also incorporates Ca^2+^ exchange between the cytoplasm and the endoplasmic reticulum (ER) through sarcoplasmic/ER Ca²⁺-ATPase (SERCA), inositol 1,4,5 trisphosphate (IP_3_) receptors, and Ca^2+^ leak channels. In addition, the Na⁺/K⁺/2Cl⁻ cotransporter 1 (NKCC1) and the K⁺/Cl⁻ cotransporter 1 (KCC1), which are integral to the regulation of ionic balance, are included in the model. NKCC1 mediates the concurrent uptake of Na⁺, K⁺, and Cl⁻ from the extracellular space, contributing to the maintenance of intracellular ion concentrations, especially during conditions of elevated extracellular K⁺. Conversely, KCC1 exports K⁺ and Cl⁻ ions from the cell, playing a crucial role in the regulation of osmotic balance and volume recovery. Together, these transporters enable astrocytes to maintain their homeostatic functions while responding to neuronal activity.

All the above considerations lead to the following rate equations for the dynamics membrane potential (*V*) and ion concentrations in the extra- and intracellular spaces. All currents are in *µM/s*, and description of all symbols and their values are given in Supplementary Table 2. The initial values of different variables are given in Supplementary Table 3.

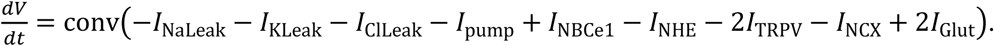

In the following, 𝑔*_pathway_* represents the maximum conductance of the respective ion pathway, and 𝐸*_ion/pathway_* represents the corresponding reversal potential.

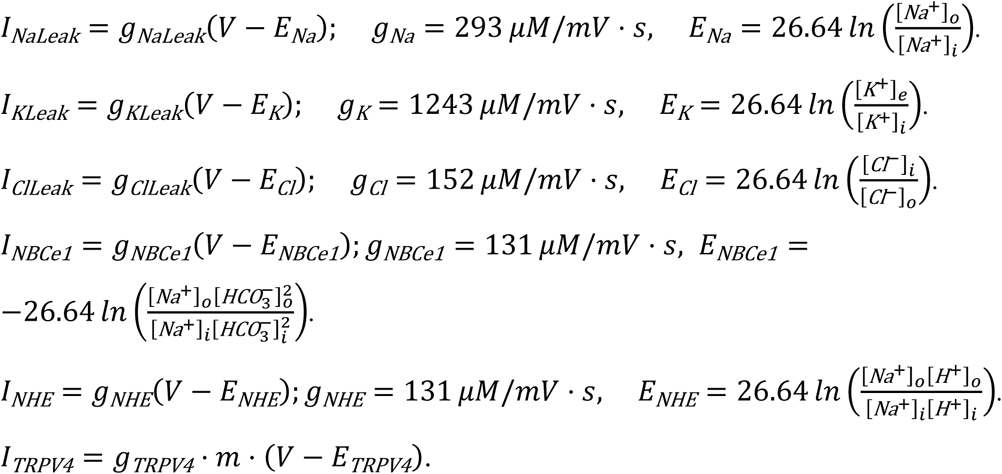

Here, the subscript *i* and *o* represent concentration in the intra- and extracellular space, respectively. Extracellular *[K^+^]* is represented by *[K^+^]_e_*. *m* is the open probability of the

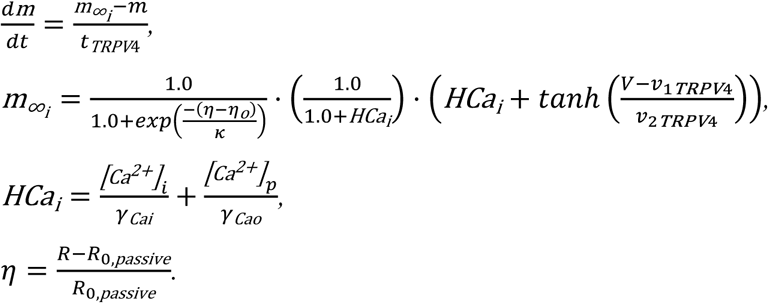

Where [*Ca*^2+^]_𝑝𝑝_ is the perivascular Ca^2+^ concentration, and the gating variable (*m*) for TRPV4 channel given by the following set of equations.

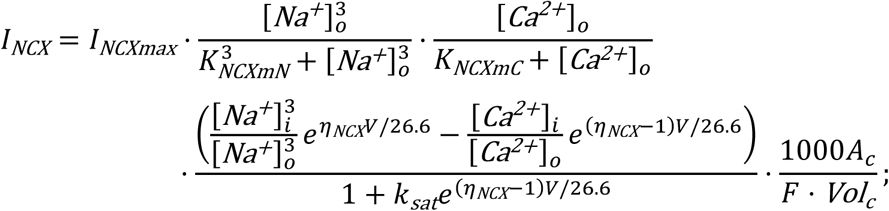

The fluxes through NCX and glutamate transporters are modeled as

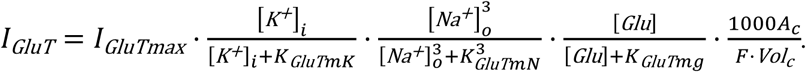

Where 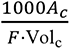 converts mS/cm^2^ to 𝜇*M*/𝑠 and 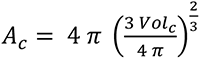 is the surface area of the cell and *Vol_c_* is its volume.

The concentrations of various ions are modeled with the following equations.

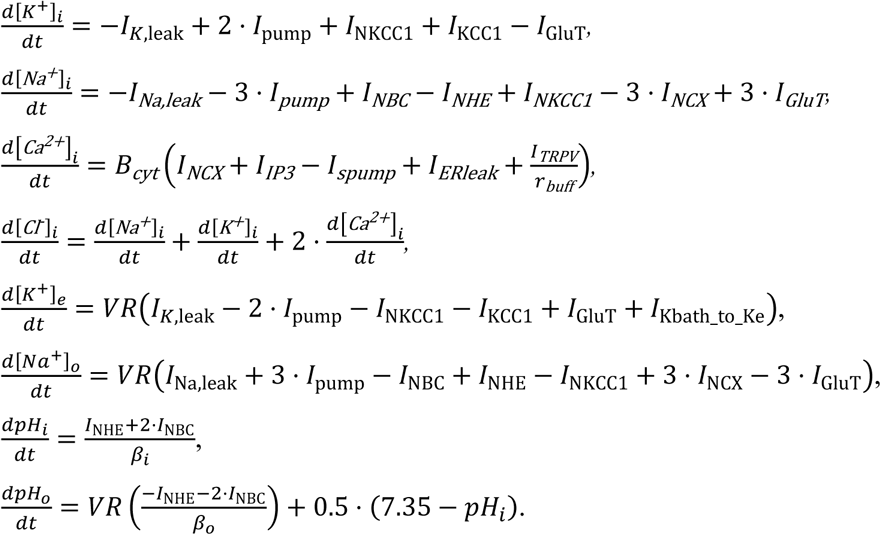

The extra- and intracellular bicarbonate and proton concentrations are given as

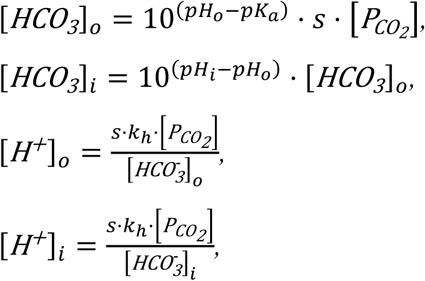

The intrinsic intra- and extracellular buffering capacity of bicarbonate and intracellular Ca^2+^ are modeled as

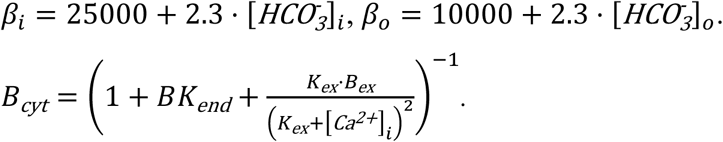

The flux through *α_2_β_1_* and *α_2_β_2_* isoforms of NKA are modeled by using the previously reported values for their binding affinities for intracellular Na^+^ and extracellular K^+^.

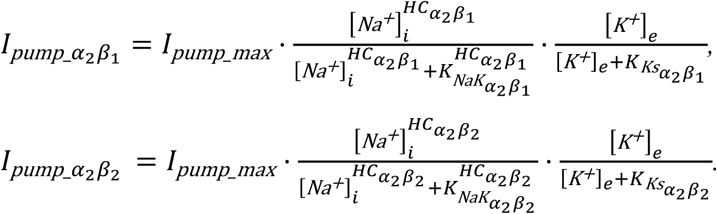

These two fluxes, normalized to their peak values, are shown in Figure S1.

The activity of NKCC1 and KCC1, and the diffusion of K^+^ from bath solution to extracellular space are given as

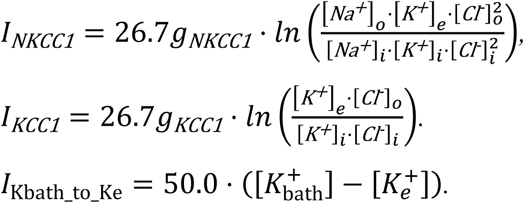

Ca^2+^ fluxes through pathways in the ER membrane (IP_3_R, SERCA, and leak) are modeled as

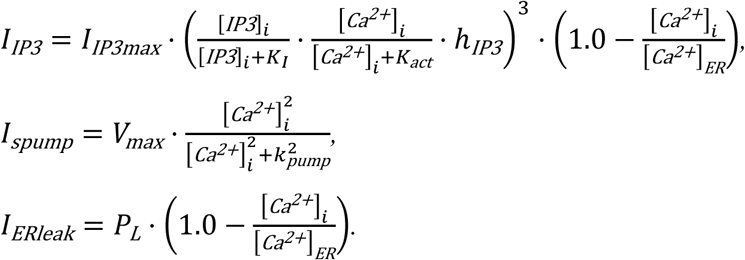

Where the concentration IP_3_ depends on the production due to glutamate (*G*) and decay over time.

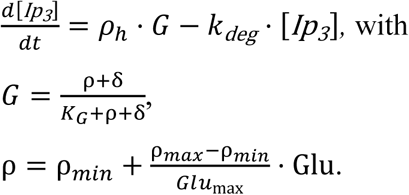

---

**Supplementary Table 2.**
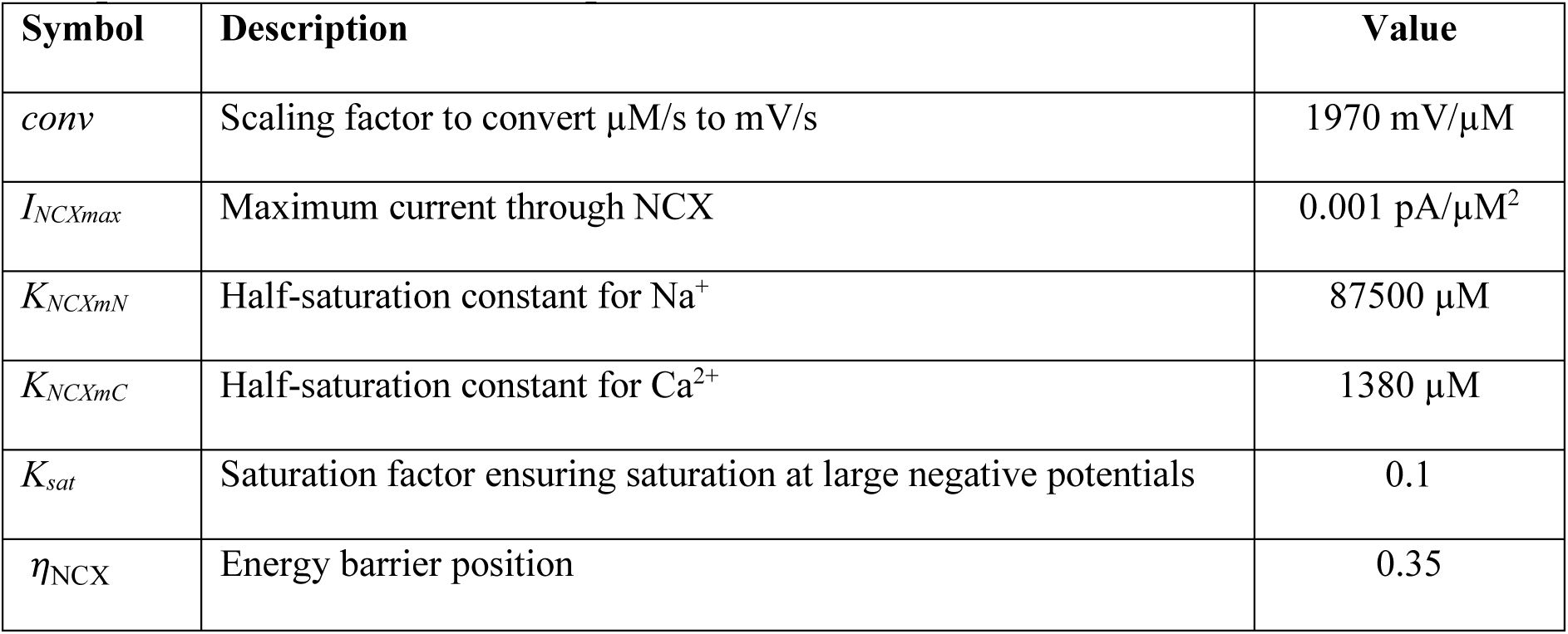
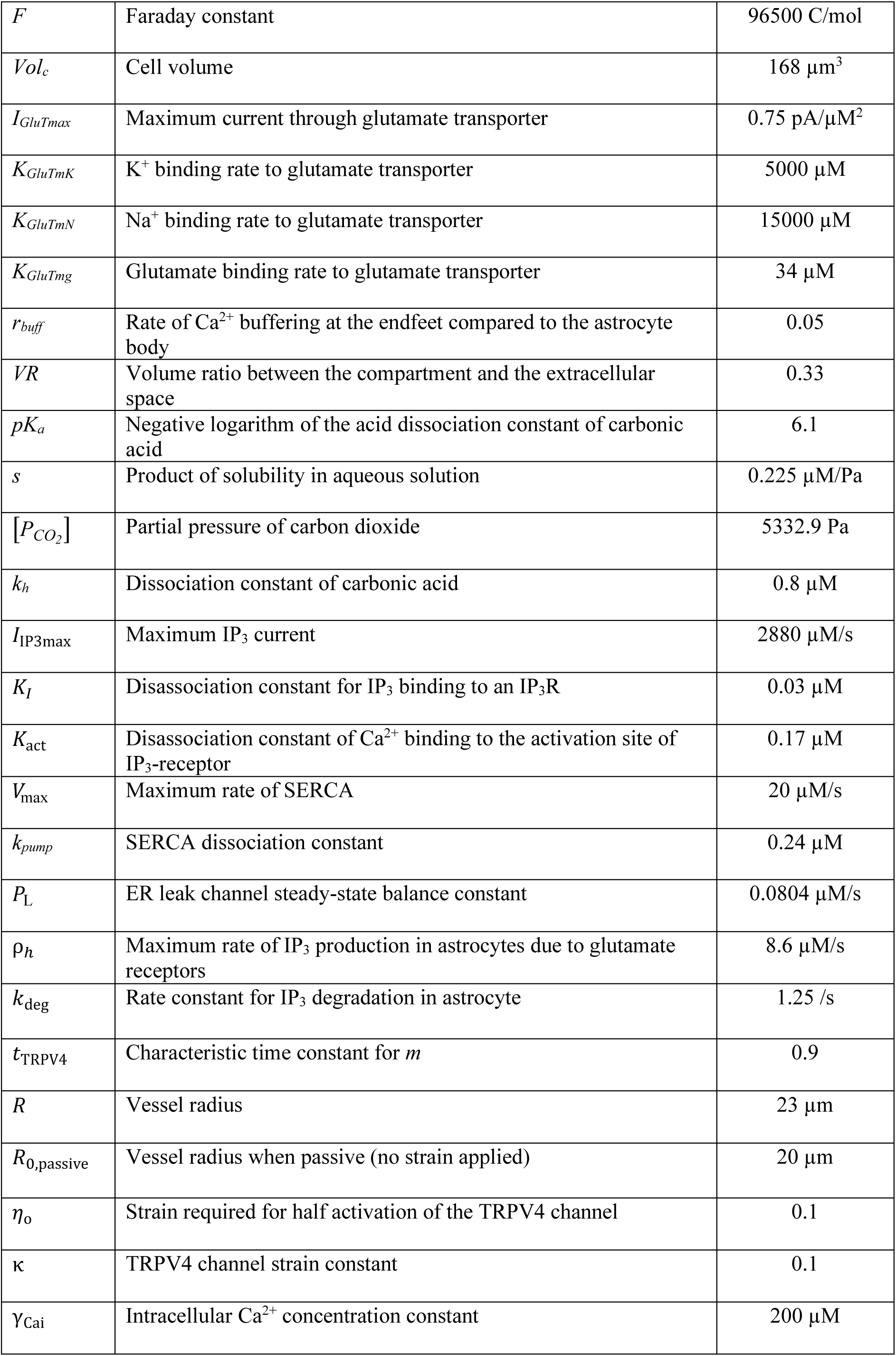
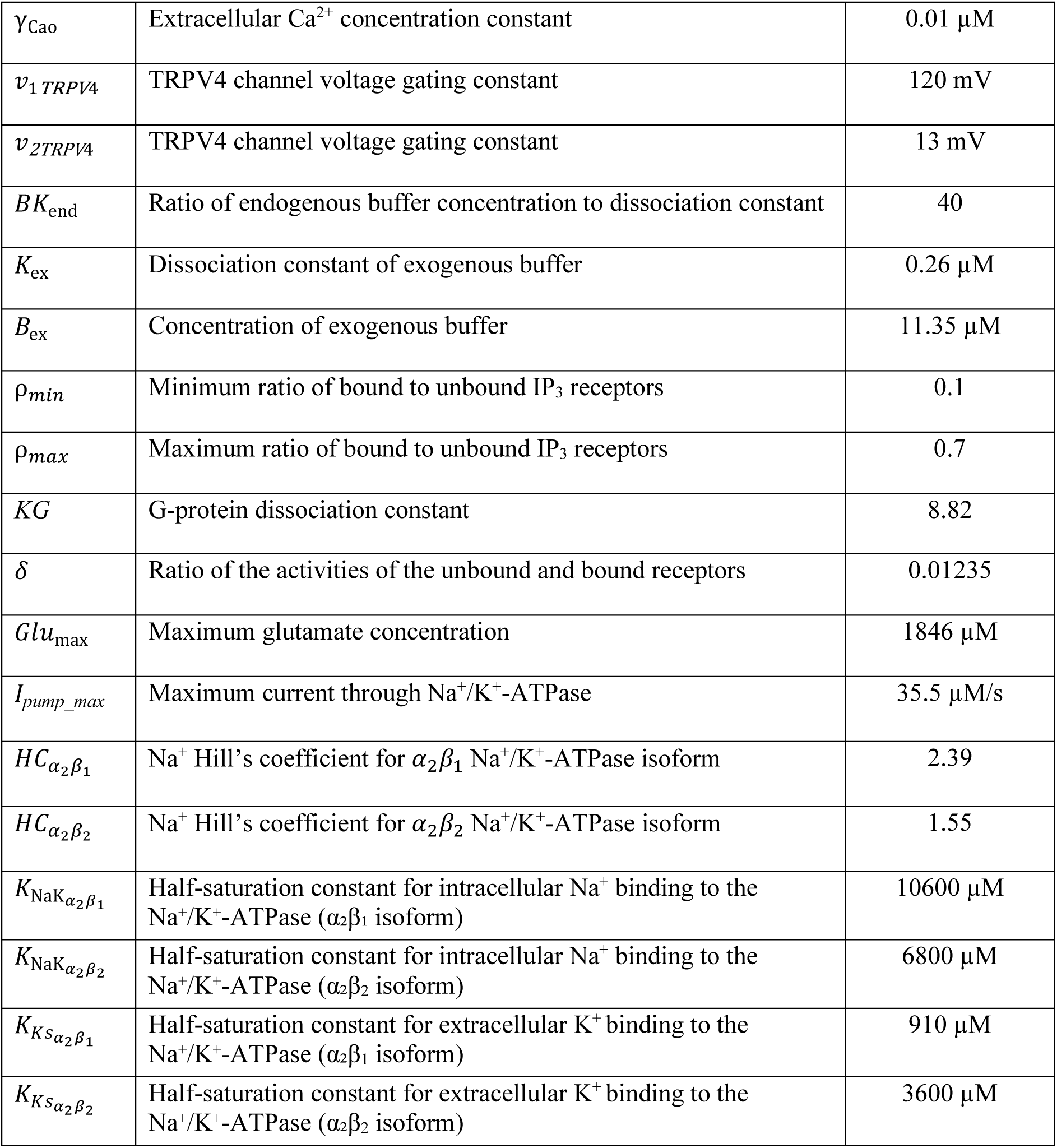
Description and values of various parameters used in the model.

| Symbol | Description | Value |
| --- | --- | --- |
| $conv$ | Scaling factor to convert $\mu\text{M/s}$ to $\text{mV/s}$ | $1970 \text{ mV}/\mu\text{M}$ |
| $I_{NCXmax}$ | Maximum current through NCX | $0.001 \text{ pA}/\mu\text{M}^2$ |
| $K_{NCXmN}$ | Half-saturation constant for $\text{Na}^+$ | $87500 \mu\text{M}$ |
| $K_{NCXmC}$ | Half-saturation constant for $\text{Ca}^{2+}$ | $1380 \mu\text{M}$ |
| $K_{sat}$ | Saturation factor ensuring saturation at large negative potentials | 0.1 |
| $\eta_{NCX}$ | Energy barrier position | 0.35 |
| $F$ | Faraday constant | 96500 C/mol |
| $Vol_c$ | Cell volume | 168 $\mu\text{m}^3$ |
| $I_{GluTmax}$ | Maximum current through glutamate transporter | 0.75 pA/$\mu\text{M}^2$ |
| $K_{GluTmK}$ | $\text{K}^+$ binding rate to glutamate transporter | 5000 $\mu\text{M}$ |
| $K_{GluTmN}$ | $\text{Na}^+$ binding rate to glutamate transporter | 15000 $\mu\text{M}$ |
| $K_{GluTmg}$ | Glutamate binding rate to glutamate transporter | 34 $\mu\text{M}$ |
| $r_{buff}$ | Rate of $\text{Ca}^{2+}$ buffering at the endfeet compared to the astrocyte body | 0.05 |
| $VR$ | Volume ratio between the compartment and the extracellular space | 0.33 |
| $pK_a$ | Negative logarithm of the acid dissociation constant of carbonic acid | 6.1 |
| $s$ | Product of solubility in aqueous solution | 0.225 $\mu\text{M}/\text{Pa}$ |
| $[P_{CO_2}]$ | Partial pressure of carbon dioxide | 5332.9 Pa |
| $k_h$ | Dissociation constant of carbonic acid | 0.8 $\mu\text{M}$ |
| $I_{IP3max}$ | Maximum $\text{IP}_3$ current | 2880 $\mu\text{M}/\text{s}$ |
| $K_I$ | Disassociation constant for $\text{IP}_3$ binding to an $\text{IP}_3\text{R}$ | 0.03 $\mu\text{M}$ |
| $K_{act}$ | Disassociation constant of $\text{Ca}^{2+}$ binding to the activation site of $\text{IP}_3$-receptor | 0.17 $\mu\text{M}$ |
| $V_{max}$ | Maximum rate of SERCA | 20 $\mu\text{M}/\text{s}$ |
| $k_{pump}$ | SERCA dissociation constant | 0.24 $\mu\text{M}$ |
| $P_L$ | ER leak channel steady-state balance constant | 0.0804 $\mu\text{M}/\text{s}$ |
| $\rho_h$ | Maximum rate of $\text{IP}_3$ production in astrocytes due to glutamate receptors | 8.6 $\mu\text{M}/\text{s}$ |
| $k_{deg}$ | Rate constant for $\text{IP}_3$ degradation in astrocyte | 1.25 /s |
| $t_{TRPV4}$ | Characteristic time constant for $m$ | 0.9 |
| $R$ | Vessel radius | 23 $\mu\text{m}$ |
| $R_{0,passive}$ | Vessel radius when passive (no strain applied) | 20 $\mu\text{m}$ |
| $\eta_o$ | Strain required for half activation of the TRPV4 channel | 0.1 |
| $\kappa$ | TRPV4 channel strain constant | 0.1 |
| $\gamma_{\text{Cai}}$ | Intracellular $\text{Ca}^{2+}$ concentration constant | 200 $\mu\text{M}$ |
| $\gamma_{CaO}$ | Extracellular $Ca^{2+}$ concentration constant | 0.01 $\mu M$ |
| $v_{1TRPV4}$ | TRPV4 channel voltage gating constant | 120 mV |
| $v_{2TRPV4}$ | TRPV4 channel voltage gating constant | 13 mV |
| $BK_{end}$ | Ratio of endogenous buffer concentration to dissociation constant | 40 |
| $K_{ex}$ | Dissociation constant of exogenous buffer | 0.26 $\mu M$ |
| $B_{ex}$ | Concentration of exogenous buffer | 11.35 $\mu M$ |
| $\rho_{min}$ | Minimum ratio of bound to unbound $IP_3$ receptors | 0.1 |
| $\rho_{max}$ | Maximum ratio of bound to unbound $IP_3$ receptors | 0.7 |
| $KG$ | G-protein dissociation constant | 8.82 |
| $\delta$ | Ratio of the activities of the unbound and bound receptors | 0.01235 |
| $Glu_{max}$ | Maximum glutamate concentration | 1846 $\mu M$ |
| $I_{pump\_max}$ | Maximum current through $Na^+/K^+$-ATPase | 35.5 $\mu M/s$ |
| $HC_{\alpha_2\beta_1}$ | $Na^+$ Hill's coefficient for $\alpha_2\beta_1$ $Na^+/K^+$-ATPase isoform | 2.39 |
| $HC_{\alpha_2\beta_2}$ | $Na^+$ Hill's coefficient for $\alpha_2\beta_2$ $Na^+/K^+$-ATPase isoform | 1.55 |
| $K_{NaK_{\alpha_2\beta_1}}$ | Half-saturation constant for intracellular $Na^+$ binding to the $Na^+/K^+$-ATPase ($\alpha_2\beta_1$ isoform) | 10600 $\mu M$ |
| $K_{NaK_{\alpha_2\beta_2}}$ | Half-saturation constant for intracellular $Na^+$ binding to the $Na^+/K^+$-ATPase ($\alpha_2\beta_2$ isoform) | 6800 $\mu M$ |
| $K_{KS_{\alpha_2\beta_1}}$ | Half-saturation constant for extracellular $K^+$ binding to the $Na^+/K^+$-ATPase ($\alpha_2\beta_1$ isoform) | 910 $\mu M$ |
| $K_{KS_{\alpha_2\beta_2}}$ | Half-saturation constant for extracellular $K^+$ binding to the $Na^+/K^+$-ATPase ($\alpha_2\beta_2$ isoform) | 3600 $\mu M$ |

**Supplementary Table 3.** Initial values of different variables

Initial values of different variables
| Variable | Value |
| --- | --- |
| $[Na^+]_i$ | 17000 μM |
| $[K^+]_i$ | 146000 μM |
| $[Ca^+]_i$ | 0.082 μM |
| $[Na^+]_o$ | 152000 μM |
| $[K^+]_e$ | 2900 μM |
| $V$ | -80 mV |
| $[IP_3]_i$ | 0.0483 μM |
| $pH_i$ | 7.33 |
| $pH_o$ | 7.35 |

**Figure S1.**
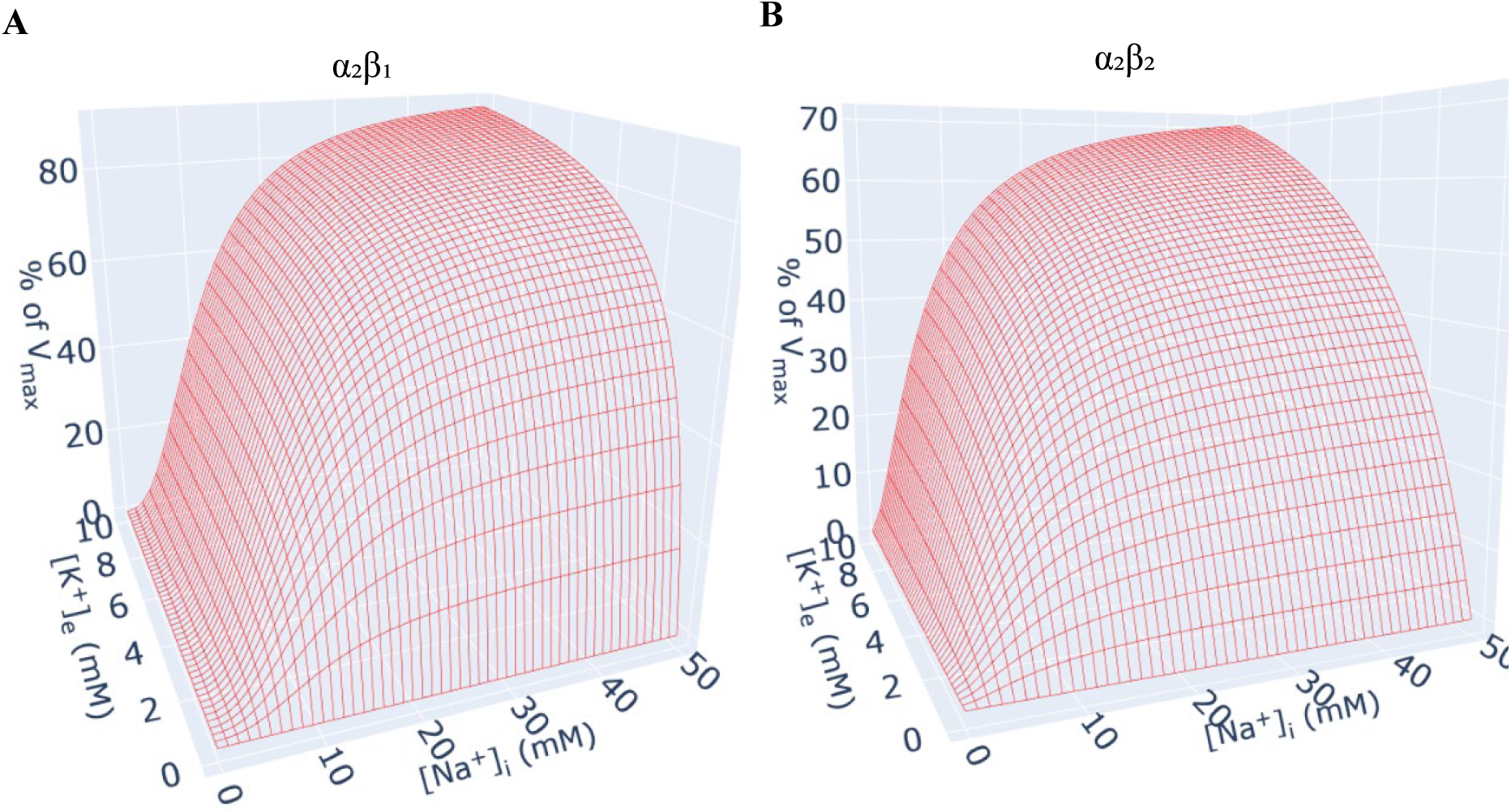
Differential activity profiles of the two isoforms of NKA as functions of intracellular Na^+^ and extracellular K^+^. The wireframe plots show the activity of NKA, normalized to their peak values (V_max_) as we change [Na^+^]_i_ and [K^+^]_e_ for α2β1 (A) and α2β2 (B). Supplementary Table 2 lists the corresponding values for Hill’s coefficients and affinities of both isoforms for [Na^+^]_i_ and [K^+^]_e_.

**Figure S2:**
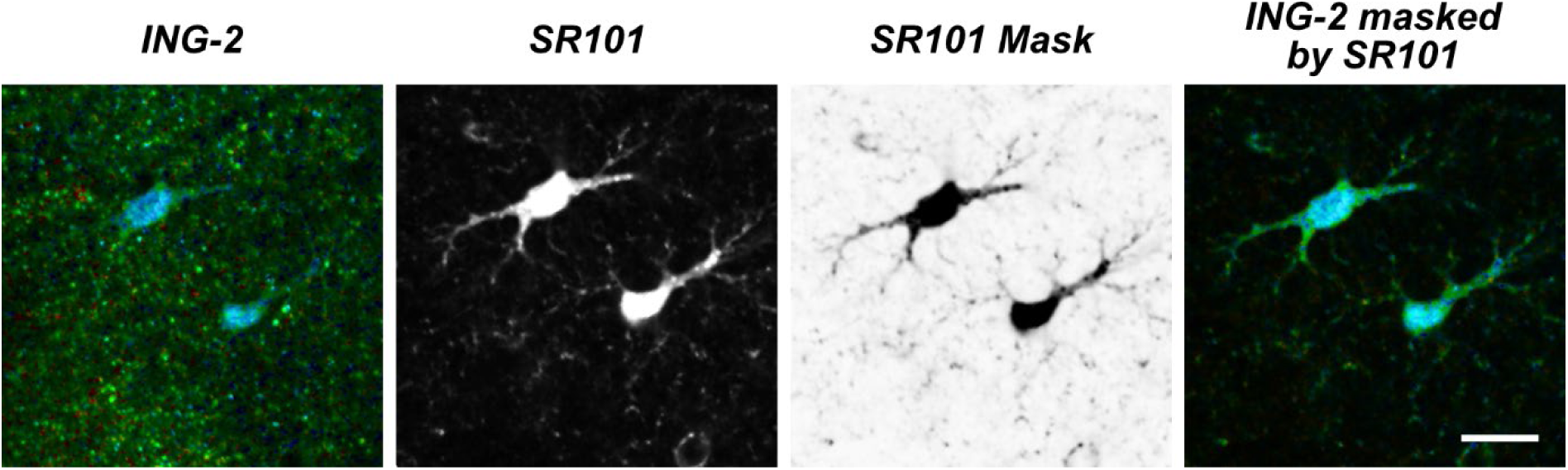
Masking ING-2 fluorescence lifetime with SR101 intensity to improve visualization of processes. From left to right: ING-2 fluorescence lifetime (FL) image, SR101 intensity image, inverted SR101 intensity image employed as a mask and ING-2 FL image masked by SR101. Utilizing the SR101 intensity image to mask the ING-2 LT image results in greatly increased visual perception of astrocyte processes. Scale: 10 µm.

## References

1. Goenaga, J., Araque, A., Kofuji, P. & Herrera Moro Chao, D. Calcium signaling in astrocytes and gliotransmitter release. Frontiers in synaptic neuroscience 15, 1138577 (2023).

2. Oliveira, J.F. & Araque, A. Astrocyte regulation of neural circuit activity and network states. Glia 70, 1455–1466 (2022).

3. Allen, N.J. & Eroglu, C. Cell Biology of Astrocyte-Synapse Interactions. Neuron 96, 697–708 (2017).

4. Verkhratsky, A. & Nedergaard, M. Physiology of Astroglia. Physiol Rev 98, 239–389 (2018).

5. Song, S., Luo, L., Sun, B. & Sun, D. Roles of glial ion transporters in brain diseases. Glia 68, 472–494 (2020).

6. Larsen, B.R. et al. Contributions of the Na(+) /K(+) -ATPase, NKCC1, and Kir4.1 to hippocampal K(+) clearance and volume responses. Glia 62, 608–622 (2014).

7. Larsen, B.R., Stoica, A. & MacAulay, N. Managing Brain Extracellular K^+^ during Neuronal Activity: The Physiological Role of the Na^+^/K^+^-ATPase Subunit Isoforms. Frontiers in physiology 7, 141 (2016).

8. Hertz, L., Song, D., Xu, J., Peng, L. & Gibbs, M.E. Role of the Astrocytic Na^+^, K^+^-ATPase in K^+^ Homeostasis in Brain: K^+^ Uptake, Signaling Pathways and Substrate Utilization. Neurochem Res (2015).

9. Tyurikova, O. et al. Astrocyte Kir4.1 expression level territorially controls excitatory transmission in the brain. Cell reports 44, 115299 (2025).

10. Wang, F. et al. Astrocytes modulate neural network activity by Ca^2+^-dependent uptake of extracellular K^+^. Sci Signal 5, ra26 (2012).

11. Aperia, A., Akkuratov, E.E., Fontana, J.M. & Brismar, H. Na+-K+-ATPase, a new class of plasma membrane receptors. Am J Physiol Cell Physiol 310, C491–495 (2016).

12. Pietrobon, D. & Conti, F. Astrocytic Na(+), K(+) ATPases in physiology and pathophysiology. Cell Calcium 118, 102851 (2024).

13. Rose, C.R. & Karus, C. Two sides of the same coin: sodium homeostasis and signaling in astrocytes under physiological and pathophysiological conditions. Glia 61, 1191–1205 (2013).

14. Rose, C.R. & Verkhratsky, A. Sodium homeostasis and signalling: The core and the hub of astrocyte function. Cell Calcium 117, 102817 (2024).

15. Boscia, F. et al. Glial Na(+) -dependent ion transporters in pathophysiological conditions. Glia 64, 1677–1697 (2016).

16. Chatton, J.Y., Magistretti, P.J. & Barros, L.F. Sodium signaling and astrocyte energy metabolism. Glia 64, 1667–1676 (2016).

17. Kirischuk, S., Heja, L., Kardos, J. & Billups, B. Astrocyte sodium signaling and the regulation of neurotransmission. Glia 64, 1655–1666 (2016).

18. Langer, J., Stephan, J., Theis, M. & Rose, C.R. Gap junctions mediate intercellular spread of sodium between hippocampal astrocytes in situ. Glia 60, 239–252 (2012).

19. Langer, J. et al. Rapid sodium signaling couples glutamate uptake to breakdown of ATP in perivascular astrocyte endfeet. Glia 65, 293–308 (2017).

20. Moshrefi-Ravasdjani, B., Hammel, E.L., Kafitz, K.W. & Rose, C.R. Astrocyte sodium signalling and panglial spread of sodium signals in brain white matter. Neurochem Res 42, 2505–2518 (2017).

21. Bernardinelli, Y., Magistretti, P.J. & Chatton, J.Y. Astrocytes generate Na^+^-mediated metabolic waves. Proc Natl Acad Sci U S A 101, 14937–14942 (2004).

22. Breslin, K. et al. Potassium and sodium microdomains in thin astroglial processes: A computational model study. PLoS Comput Biol 14, e1006151 (2018).

23. Wade, J.J. et al. Calcium Microdomain Formation at the Perisynaptic Cradle Due to NCX Reversal: A Computational Study. Frontiers in cellular neuroscience 13, 185 (2019).

24. Armbruster, M. et al. Neuronal activity drives pathway-specific depolarization of peripheral astrocyte processes. Nat Neurosci 25, 607–616 (2022).

25. Nakatani, R.J. & De Schutter, E. Active enhancement of synapse driven depolarization of perisynaptic astrocytic processes. bioRxiv, 2024.2006.2005.597669 (2024).

26. Herde, M.K. et al. Local Efficacy of Glutamate Uptake Decreases with Synapse Size. Cell reports 32, 108182 (2020).

27. Henneberger, C. et al. LTP Induction Boosts Glutamate Spillover by Driving Withdrawal of Perisynaptic Astroglia. Neuron 108, 919–936 e911 (2020).

28. Becker, W. Advanced Time-Correlated Single Photon Counting Applications. (Springer-Verlag, Berlin Heidelberg, 2015).

29. Kuchibhotla, K.V., Lattarulo, C.R., Hyman, B.T. & Bacskai, B.J. Synchronous hyperactivity and intercellular calcium waves in astrocytes in Alzheimer mice. Science 323, 1211–1215 (2009).

30. Untiet, V. et al. Glutamate transporter-associated anion channels adjust intracellular chloride concentrations during glial maturation. Glia 65, 388–400 (2017).

31. Zheng, K. et al. Time-Resolved Imaging Reveals Heterogeneous Landscapes of Nanomolar Ca(2+) in Neurons and Astroglia. Neuron 88, 277–288 (2015).

32. King, C.M. et al. Local Resting Ca(2+) Controls the Scale of Astroglial Ca(2+) Signals. Cell reports 30, 3466–3477 e3464 (2020).

33. Meyer, J. et al. Rapid fluorescence lifetime imaging reveals that TRPV4 channels promote dysregulation of neuronal Na(+) in ischemia. J Neurosci 42, 552–566 (2022).

34. Nelson, J.S.E. et al. Spatio-temporal dynamics of lateral Na+ diffusion in apical dendrites of mouse CA1 pyramidal neurons. bioRxiv, 2025.2008.2006.668873 (2025).

35. Batiuk, M.Y. et al. Identification of region-specific astrocyte subtypes at single cell resolution. Nature communications 11, 1220 (2020).

36. Bayraktar, O.A. et al. Astrocyte layers in the mammalian cerebral cortex revealed by a single-cell in situ transcriptomic map. Nat Neurosci 23, 500–509 (2020).

37. Chai, H. et al. Neural Circuit-Specialized Astrocytes: Transcriptomic, Proteomic, Morphological, and Functional Evidence. Neuron 95, 531–549 e539 (2017).

38. de Ceglia, R. et al. Specialized astrocytes mediate glutamatergic gliotransmission in the CNS. Nature 622, 120–129 (2023).

39. Dossi, E. et al. Astroglial gap junctions strengthen hippocampal network activity by sustaining afterhyperpolarization via KCNQ channels. Cell reports 43, 114158 (2024).

40. Karus, C., Mondragao, M.A., Ziemens, D. & Rose, C.R. Astrocytes restrict discharge duration and neuronal sodium loads during recurrent network activity. Glia 63, 936–957 (2015).

41. Clarke, L.E. et al. Normal aging induces A1-like astrocyte reactivity. Proc Natl Acad Sci U S A 115, E1896–E1905 (2018).

42. Zhang, Y. et al. An RNA-Sequencing Transcriptome and Splicing Database of Glia, Neurons, and Vascular Cells of the Cerebral Cortex. J Neurosci 34, 11929–11947 (2014).

43. Bernardinelli, Y., Azarias, G. & Chatton, J.Y. In situ fluorescence imaging of glutamate-evoked mitochondrial Na^+^ responses in astrocytes. Glia 54, 460–470 (2006).

44. Unichenko, P., Myakhar, O. & Kirischuk, S. Intracellular Na(+) concentration influences short-term plasticity of glutamate transporter-mediated currents in neocortical astrocytes. Glia 60, 605–614 (2012).

45. Langer, J. & Rose, C.R. Synaptically induced sodium signals in hippocampal astrocytes in situ. J Physiol 587, 5859–5877 (2009).

46. Ziemens, D., Oschmann, F., Gerkau, N.J. & Rose, C.R. Heterogeneity of activity-induced sodium transients between astrocytes of the mouse hippocampus and neocortex: Mechanisms and consequences. J Neurosci 39, 2620–2634 (2019).

47. Theparambil, S.M. et al. Bicarbonate sensing in mouse cortical astrocytes during extracellular acid/base disturbances. J Physiol 595, 2569–2585 (2017).

48. Noor, Z.N., Deitmer, J.W. & Theparambil, S.M. Cytosolic sodium regulation in mouse cortical astrocytes and its dependence on potassium and bicarbonate. J Cell Physiol 234, 89–99 (2018).

49. Rose, C.R. & Ransom, B.R. Gap junctions equalize intracellular Na^+^ concentration in astrocytes. Glia 20, 299–307 (1997).

50. Pannasch, U. et al. Astroglial networks scale synaptic activity and plasticity. Proc Natl Acad Sci U S A 108, 8467–8472 (2011).

51. Rouach, N., Segal, M., Koulakoff, A., Giaume, C. & Avignone, E. Carbenoxolone blockade of neuronal network activity in culture is not mediated by an action on gap junctions. J Physiol 553, 729–745 (2003).

52. Tovar, K.R., Maher, B.J. & Westbrook, G.L. Direct actions of carbenoxolone on synaptic transmission and neuronal membrane properties. J Neurophysiol 102, 974–978 (2009).

53. Clarke, B.E., Taha, D.M., Tyzack, G.E. & Patani, R. Regionally encoded functional heterogeneity of astrocytes in health and disease: A perspective. Glia 69, 20–27 (2021).

54. Hasel, P., Aisenberg, W.H., Bennett, F.C. & Liddelow, S.A. Molecular and metabolic heterogeneity of astrocytes and microglia. Cell metabolism 35, 555–570 (2023).

55. Lanjakornsiripan, D. et al. Layer-specific morphological and molecular differences in neocortical astrocytes and their dependence on neuronal layers. Nature communications 9, 1623 (2018).

56. Stoica, A. et al. The alpha2beta2 isoform combination dominates the astrocytic Na+ /K+ - ATPase activity and is rendered nonfunctional by the alpha2.G301R familial hemiplegic migraine type 2-associated mutation. Glia 65, 1777–1793 (2017).

57. Cholet, N., Pellerin, L., Magistretti, P.J. & Hamel, E. Similar perisynaptic glial localization for the Na+,K+-ATPase alpha 2 subunit and the glutamate transporters GLAST and GLT-1 in the rat somatosensory cortex. Cereb Cortex 12, 515–525 (2002).

58. Melone, M., Ciriachi, C., Pietrobon, D. & Conti, F. Heterogeneity of Astrocytic and Neuronal GLT-1 at Cortical Excitatory Synapses, as Revealed by its Colocalization With Na+/K+-ATPase alpha Isoforms. Cereb Cortex 29, 3331–3350 (2019).

59. Lehre, K.P. & Danbolt, N.C. The number of glutamate transporter subtype molecules at glutamatergic synapses: chemical and stereological quantification in young adult rat brain. J Neurosci 18, 8751–8757 (1998).

60. Chaudhry, F.A. et al. Glutamate transporters in glial plasma membranes: highly differentiated localizations revealed by quantitative ultrastructural immunocytochemistry. Neuron 15, 711–720 (1995).

61. Radulescu, A.R. et al. Estimating the glutamate transporter surface density in distinct sub-cellular compartments of mouse hippocampal astrocytes. PLoS Comput Biol 18, e1009845 (2022).

62. Illarionava, N.B., Brismar, H., Aperia, A. & Gunnarson, E. Role of Na, K-ATPase alpha1 and alpha2 isoforms in the support of astrocyte glutamate uptake. PLoS One 9, e98469 (2014).

63. Larsen, B.R., Holm, R., Vilsen, B. & MacAulay, N. Glutamate transporter activity promotes enhanced Na+ /K+ -ATPase-mediated extracellular K+ management during neuronal activity. J Physiol 594, 6627–6641 (2016).

64. Meyer, J., Kafitz, K.W. & Rose, C.R. Quantification of Astrocytic Sodium Signals Using Fluorescence Lifetime Imaging Microscopy (FLIM). Methods Mol Biol 2896, 51–61 (2025).

65. Close, B. et al. Recommendations for euthanasia of experimental animals: Part 2. DGXT of the European Commission. Laboratory animals 31, 1–32 (1997).

66. Rose, C.R. & Ransom, B.R. Intracellular sodium homeostasis in rat hippocampal astrocytes. J Physiol 491, 291–305 (1996).

67. Gerkau, N.J., Rakers, C., Durry, S., Petzold, G.C. & Rose, C.R. Reverse NCX Attenuates Cellular Sodium Loading in Metabolically Compromised Cortex. Cereb Cortex 28, 4264–4280 (2018).

68. Minge, D. et al. Heterogeneity and Development of Fine Astrocyte Morphology Captured by Diffraction-Limited Microscopy. Frontiers in cellular neuroscience 15, 669280 (2021).

69. Haack, N., Durry, S., Kafitz, K.W., Chesler, M. & Rose, C.R. Double-barreled and concentric microelectrodes for measurement of extracellular ion signals in brain tissue. Journal of visualized experiments : JoVE, doi: 10.3791/53058 (2015).

## References

1. Everaerts K, Thapaliya P, Pape N, Durry S, Eitelmann S, Roussa E, Ullah G, Rose CR ((2023) Inward Operation of Sodium-Bicarbonate Cotransporter 1 Promotes Astrocytic Na(+) Loading and Loss of ATP in Mouse Neocortex during Brief Chemical Ischemia. Cells 12(23).

2. Thapaliya P, Pape N, Ullah G ((2023) Modeling the heterogeneity of sodium and calcium homeostasis between cortical and hippocampal astrocytes and its impact on bioenergetics. Front Cell Neurosci 17: p. 1035553.

3. Kenny A, Plank MJ, David T (2018) The role of astrocytic calcium and TRPV4 channels in neurovascular coupling. J Comput Neurosci 44(1): p. 97–114.

73. Oschmann F (2018) Computational Modeling of Glutamate-Induced Calcium Signal Generation and Propagation in Astrocytes. Dissertation, Technische Universitaet Berlin. Germany.

5. Larsen BR, Stoica A, MacAulay N (2016a) Managing Brain Extracellular K^+^ during Neuronal Activity: The Physiological Role of the Na^+^/K^+^-ATPase Subunit Isoforms. Frontiers in physiology 7:141.

6. Larsen BR, Holm R, Vilsen B, MacAulay N (2016b) Glutamate transporter activity promotes enhanced Na+ /K+ -ATPase-mediated extracellular K+ management during neuronal activity. J Physiol 594:6627–6641.

7. Zhang Y, Chen K, Sloan SA, Bennett ML, Scholze AR, O’Keeffe S, Phatnani HP, Guarnieri P, Caneda C, Ruderisch N, Deng S, Liddelow SA, Zhang C, Daneman R, Maniatis T, Barres BA, Wu JQ (2014) An RNA-Sequencing Transcriptome and Splicing Database of Glia, Neurons, and Vascular Cells of the Cerebral Cortex. J Neurosci 34:11929–11947.

8. Clarke LE, Liddelow SA, Chakraborty C, Munch AE, Heiman M, Barres BA (2018) Normal aging induces A1-like astrocyte reactivity. Proc Natl Acad Sci U S A 115:E1896–E1905.

78. Batiuk MY, Martirosyan A, Wahis J, de Vin F, Marneffe C, Kusserow C, Koeppen J, Viana JF, Oliveira JF, Voet T, Ponting CP, Belgard TG, Holt MG (2020) Identification of region-specific astrocyte subtypes at single cell resolution. Nature communications 11:1220.

